# Dorsolateral striatal acetylcholine reorganizes neural ensembles to anticipate threat

**DOI:** 10.64898/2025.12.10.693585

**Authors:** Oyku Dinckol, Noah Harris Wenger, Charlie Maddox, Taylor Good, Aryanna Copling, Bhumi Pradipkumar Patel, F. Zehra Bozdag, Munir Gunes Kutlu

**Affiliations:** Center for Substance Abuse Research (CSAR), Temple University Lewis Katz School of Medicine, Philadelphia, PA, USA; Department of Cell Biology & Neuroscience, Rowan-Virtua University School of Osteopathic Medicine, Stratford, NJ; Biomedical Sciences Graduate Program, Temple University Lewis Katz School of Medicine, Philadelphia, PA, USA; Department of Neurosciences, Lerner Research Institute, Cleveland Clinic, Cleveland, OH; Department of Neural Sciences, Temple University Lewis Katz School of Medicine, Philadelphia, PA, USA

**Keywords:** Striatum, Acetylcholine, threat learning, neural ensembles

## Abstract

Adaptive behavior requires flexible encoding of emotional valence. Although striatal acetylcholine (ACh) signaling is critical for reinforcement learning, its contribution to aversive learning has remained poorly defined. Here, we demonstrate that ACh release in the dorsolateral striatum (DLS) is selectively biased toward negative valence. Using fiber photometry with a genetically encoded ACh sensor, we found that ACh release robustly increased during threat prediction but decreased in anticipation of rewarding outcomes, revealing a bidirectional and valence-specific signature. Optogenetically stimulating ACh release at cue onset accelerated threat learning, impaired extinction, and shifted behavioral responding toward persistent threat expectancy. Concurrent single-cell calcium imaging and optogenetic manipulation revealed that elevated ACh release dynamically reorganized DLS ensemble activity, increasing both excited and inhibited neurons and producing large-scale state-space divergence during threat cues. During extinction, optogenetically sustained ACh release preserved the organization of threat-predictive DLS ensemble activity despite the absence of shock. These findings identify DLS ACh as a valence-specific neuromodulatory signal that reconfigures striatal network dynamics, primes ensembles for impending threat, and biases learning toward threat persistence.

## Introduction

Cholinergic signaling, mediated by acetylcholine (ACh), broadly contributes to neural plasticity^1,2^ and supports diverse cognitive operations, including attention/working memory^3–7^, motivation^8,9^, prediction error^10,11^, and saliency encoding^12–14^, underscoring its role in adaptive neural computation. ACh has traditionally been studied within cortical and hippocampal circuits, where its mechanisms and behavioral functions are best characterized^4,15^. However, ACh is also highly influential in subcortical regions, particularly within the striatum^16^. Nevertheless, its contribution to striatal information processing, especially its role in aversive learning, remains poorly defined.

Within the striatum, cholinergic interneurons (CINs) constitute a small but highly influential population that provides the primary source of striatal ACh^16–18^ and exerts strong modulatory control over microcircuit activity, reinforcement learning, and dopamine–GABA interactions^19–21^. Striatal cholinergic signaling has been implicated in movement initiation^7,22,23^, encoding salience^14^, and reward learning^8,24–26^. Yet, while these studies highlight the importance of striatal ACh release and CIN activity patterns in reward-related processes, their roles outside of reward learning remain largely unexplored.

Despite the central role of CINs across the striatum, the mechanisms by which cholinergic signaling contributes to aversive processing within specific striatal subregions remain unclear. The present study focuses on the dorsal striatum, a region traditionally known for its role in reward learning^27–30^ and increasingly recognized for its involvement in aversive learning^31,32^. Specifically, the dorsolateral striatum (DLS), traditionally associated with sensorimotor integration^33^ and habitual behavior^27,34,35^, has received little attention in the context of affective signaling.

Here, we ask whether ACh release in the DLS is selective for reward learning, extends similarly to aversive contexts, or instead reflects a valence-independent learning signal. We further test the causal contribution of DLS cholinergic activity to threat perception and aversive learning, a question that has not been directly examined. Finally, we determine how ACh release shapes and coordinates DLS ensemble dynamics during exposure to aversive stimuli. Addressing these questions is essential for understanding how striatal circuits integrate emotional, motivational, and motor information to guide adaptive and flexible behavior.

## Results

### DLS ACh release is selectively biased toward negatively valenced experiences

We first characterized ACh release dynamics in the DLS in response to novel stimuli of varying valence: neutral (tone), positive (30% sucrose), or negative (footshocks or quinine). Using fiber photometry in awake, behaving mice expressing the genetically encoded ACh sensor iACh.Sn.FR (AAV1.hSynap.iACh.Sn.FR^36^, see **Extended Data Fig. 1** for the validation of our ACh recordings), we recorded real-time fluctuations in extracellular ACh levels. Our results showed that the presentation of a neutral stimulus elicited a minimal ACh release response (**Extended Data Fig. 2a,b**). Interestingly, free access to 30% sucrose (a positively valenced stimulus) resulted in decreased ACh release during licks, whereas free quinine access (a negatively valenced stimulus) increased ACh release in the DLS (**Extended Data Fig. 2c,d**). Footshocks (1 mA), another negatively valenced stimulus, also robustly increased ACh release, in a manner similar to quinine, with a more dramatic increase (**Extended Data Fig. 2e,f**). These results indicate that DLS ACh release is selectively biased toward negatively valenced experiences, suggesting a valence-specific neuromodulatory role in aversion-related processing.

### DLS ACh release scales with threat learning and decreases during reward learning

Our initial findings showed that non-contingent aversive experiences increased DLS ACh release, whereas exposure to rewards suppressed it, suggesting that aversive and rewarding associative learning may bidirectionally modulate DLS ACh. To investigate this, we recorded ACh release dynamics during fear conditioning and extinction sessions (see **Fig. 1a,b** for a schematic description and **Fig. 1c** for the freezing responses). During fear conditioning, DLS ACh release increased following CS+ presentations, with a larger surge at the time of footshock delivery (**Fig. 1d,e**). Freezing during CS+ presentations positively correlated with the ACh responses during these trials (**Extended Data Fig. 3**). During extinction, when footshocks were omitted following the CS+ presentations, both freezing (**Fig. 1f**) and ACh responses to the CS+ gradually declined across sessions, with a significant reduction by extinction session 5 (EXT5) compared to extinction session 1 (EXT1, **Fig. 1g-i**). Thus, DLS ACh tracks the strength of threat-associated learning.

**Figure 1.**
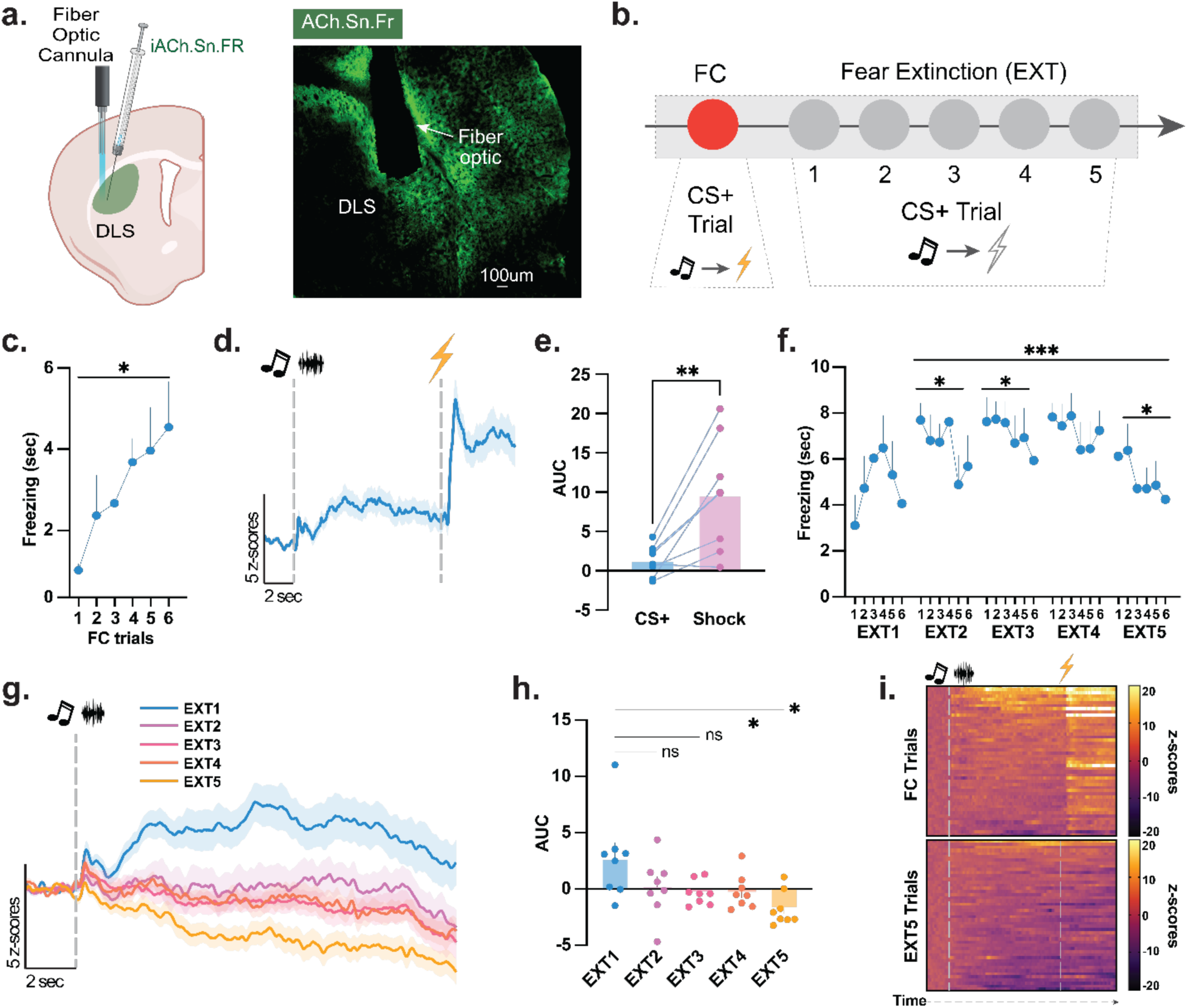
DLS ACh release tracks threat anticipation and shock delivery during fear learning. ***(a)*** Mice (n=8 mice, 5 males and 3 females) received a unilateral injection of AAV1.hSynap.iACh.Sn.FR and 400µm fiber optic implantation into the DLS to induce GFP activation when ACh binds to its receptor. Representative histology shows viral expression and ACh-dependent GFP activation (green) at synapses within the DLS, as well as validated fiber optic placement for fiber photometry recordings. ***(b)*** Mice underwent a single fear conditioning session (FC) followed by five consecutive fear extinction (EXT) sessions. During fear conditioning, animals received six CS+ presentations (85 dB tone or white noise, counterbalanced; 10 s) predicting a footshock (1 mA, 0.5 sec). During the five extinction sessions, mice received the same six CS+ presentations per session, with footshocks omitted. ***(c)*** Freezing behavior during the 10-s CS+ increased progressively across the six fear-conditioning trials, reaching robust levels by the final acquisition trial (Paired t-test [Trial 1 vs Trial 6 freezing]; *t*(7)=2.474, *p*=0.0426; n=8 mice). ***(d)*** Fiber photometry recordings revealed a robust increase in DLS ACh during each CS+ as animals anticipated the shock, followed by a rapid, larger ACh increase at the moment of shock delivery. ***(e)*** The footshock induced a significantly greater ACh release compared to the CS+ period (Paired t-test; *t*(7)=3.908, *p*=0.0058; n=8 mice). ***(f)*** During fear extinction, the freezing response to the CS+ showed a gradual decrease. Extinction training was analyzed separately across the five sessions (Twelve extinction trials were averaged in bins of two trials per session). A linear trend analysis revealed significant within-session extinction on Sessions 2 (EXT2; *p*=0.022), Session 3 (EXT3; *p*=0.058), and Session 5 (EXT5; *p*=0.027), with progressive decreases in freezing across Sessions 2 through 5 (*p*=0.0005). ***(g-h)*** During extinction, CS-evoked ACh responses progressively decreased across sessions, with significant reductions from EXT1 to EXT4 and EXT1 to EXT5 (RM ANOVA; F(2.407,16.85) = 7.327, *p*=0.0036; n=8 mice). ***(i)*** Heatmaps illustrate the average population response aligned to CS onset and shock. ACh signals were strongly elevated during conditioning (FC, top) but returned to near baseline by extinction session 5 (EXT5, bottom), consistent with reduced threat expectancy. Data represented as mean ± S.E.M. *\** p < 0.05, ** p < 0.01, *** p < 0.001.

Next, we assessed responses to a positively valenced experience and tested our animals on appetitive conditioning. Animals received cue presentations that signaled free sucrose access (CS+; tone or white noise) for 10 seconds (see **Fig. 2a** for a schematic description). Despite successful learning, animals showed successful acquisition of the conditioning, as the approach ratio significantly increased from Session 1 to Session 6 (**Fig. 2b**). DLS ACh release elicited a decreased response to CS+ and remained below baseline across learning sessions, in contrast to the increased approach behavior (**Fig. 2c,d**). These results suggest that DLS ACh release is biased towards negative valence and signals upcoming threats such as footshocks.

**Figure 2.**
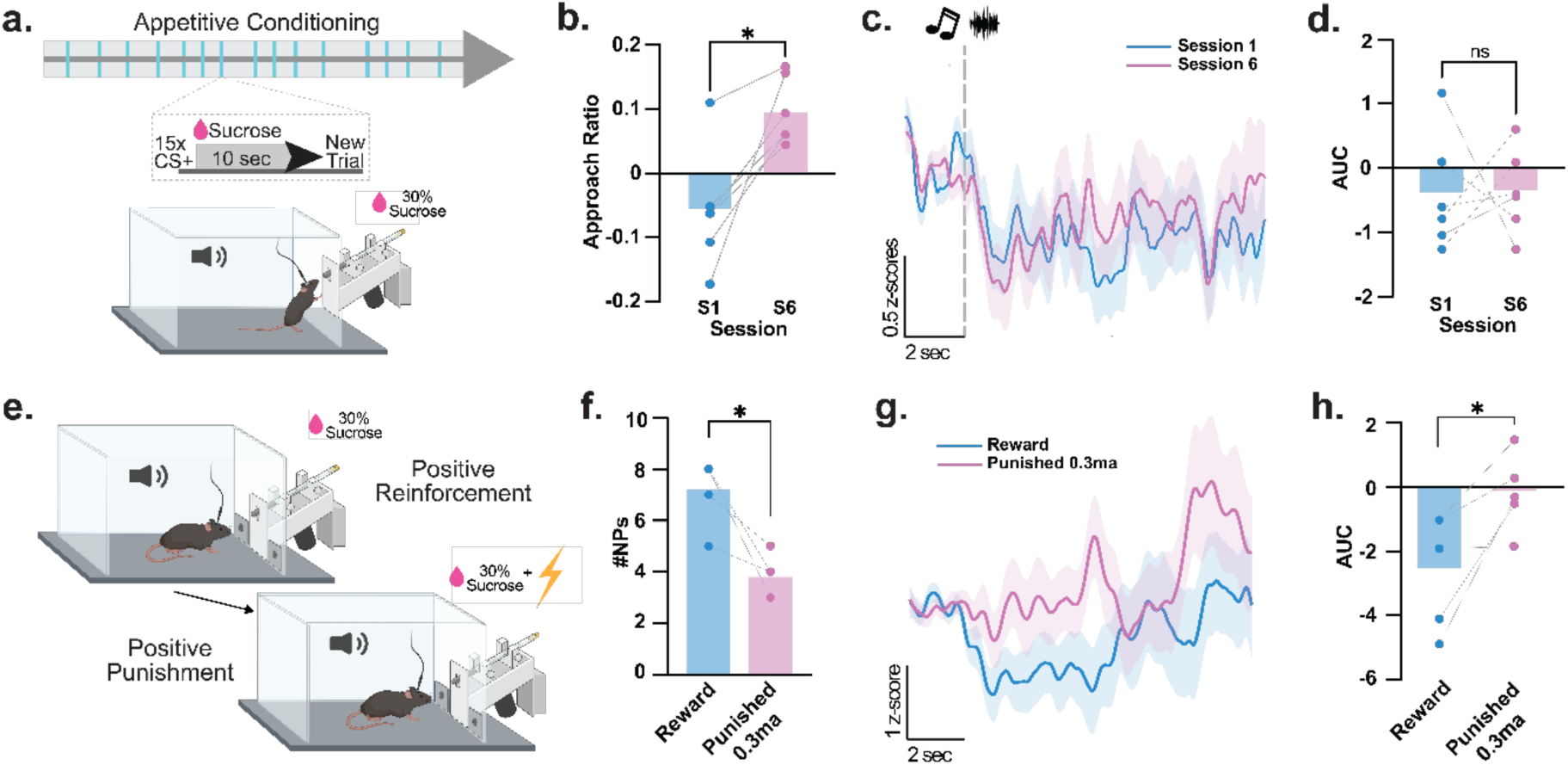
DLS ACh release bidirectionally signals prediction of reward and punishment. ***(a)*** Mice (n=6 mice, 3 males and 3 females) underwent six consecutive sessions of appetitive conditioning. Each session contained fifteen CS+ presentations (85 dB tone or white noise, counterbalanced; 10 s) that predicted access to a retractable sipper delivering sucrose (10 s). ***(b)*** Reward learning was reflected in a progressive increase in sipper approaches across sessions in the learner mice, with a significant increase in approach ratio in session 6 compared to session 1 (Paired t-test; *t*(5)=4.026, *p*=0.0101; n=6 mice). ***(c)*** Fiber photometry recordings show DLS ACh responses aligned to CS+ onset during appetitive conditioning sessions 1 and 6. ***(d)*** Quantification revealed no significant difference in CS-evoked ACh responses between sessions 1 and 6 (Paired t-test; *t*(5)=0.066, *p*=0.9494; n=6 mice). ***(e)*** For the positive reinforcement/positive punishment test, mice (n=5 mice, 3 males and 2 females) were trained on a cued FR1 task in which a correct nose poke during the auditory cue yielded access to 30% sucrose. After achieving ≥50% cue-correct responses, mice were transitioned to a cued FR1 punishment task in which obtaining sucrose resulted in a 0.3 mA, 0.5 s footshock. ***(f)*** Introduction of punishment significantly reduced correct responding (Paired t-test; *t*(4)=4.185, *p*=0.0139). ***(g)*** DLS ACh activity during reward and punishment trials. ***(h)*** Quantified AUCs show that DLS ACh release was significantly elevated during punishment sessions compared to reward sessions (Paired t-test; *t*(4)=3.016, *p*=0.0393; n=5 mice), demonstrating a bidirectional cholinergic signal reflecting outcome valence. Data represented as mean ± S.E.M. *\** p < 0.05.

Finally, we tested whether DLS ACh selectively signals negative outcomes. We predicted that ACh release patterns would invert, shifting from positive to negative, when cues or action valence transitioned from rewarding to aversive, and tested this using an operant reward-punishment paradigm (see **Fig. 2e** for a schematic description). Animals were trained to nose poke on the left side port (active nose poke) during a cue (CS+) to receive 10 seconds of 30% sucrose access. Following a training period, ACh release was measured while the animals performed the task during the reward session. Animals that acquired the task successfully (**Fig. 2f**), *learners*, had decreased ACh release (**Fig. 2g**), while animals that performed below the 50% correct response threshold, *non-learners*, had increased ACh release (**Extended Data Fig. 4**). Following the reward session, learners proceeded to the punishment session, where performing an active nose-poke delivered both a footshock and sucrose access. Animals significantly reduced active nose-poking during this session (**Fig. 2f**), and the ACh release response to the CS+ increased (**Fig. 2g,h**), suggesting that DLS ACh response scales with threat perception during a predictive cue. Together, the results across fear conditioning/extinction and appetitive conditioning paradigms indicate that DLS ACh reflects threat perception rather than reward, dynamically tracking aversive learning and predictive cues.

### DLS ACh causally mediates threat perception during fear learning

Next, we examined whether the DLS ACh release causally contributes to threat perception by optogenetically stimulating local CINs. We injected an AAV2/5-ChAT-Cre-WPRE-HGH virus to drive Cre-recombinase expression in CINs and expressed either a Cre-dependent excitatory opsin (AAV5-Syn-FLEX-rc[ChrimsonR-tdTomato]; *Chrimson* animals) for optogenetic excitation or a control virus (AAV5-hSyn-DIO-mCherry; *mCherry* animals) (**Fig. 3a**). Cre-dependent Chrimson expression in ChAT neurons was verified histologically in both the basal forebrain and DLS (**Supp Fig. 5**). All optogenetic animals also received iACh.Sn.FR (AAV1.hSynap.iACh.Sn.FR; **Fig. 3a**) to validate DLS ACh release during laser stimulation. Optogenetic excitation of CINs at different laser intensities (10Hz 2mW, 5mW, 8mW) reliably increased ACh release (**Fig. 3b,c**).

**Figure 3.**
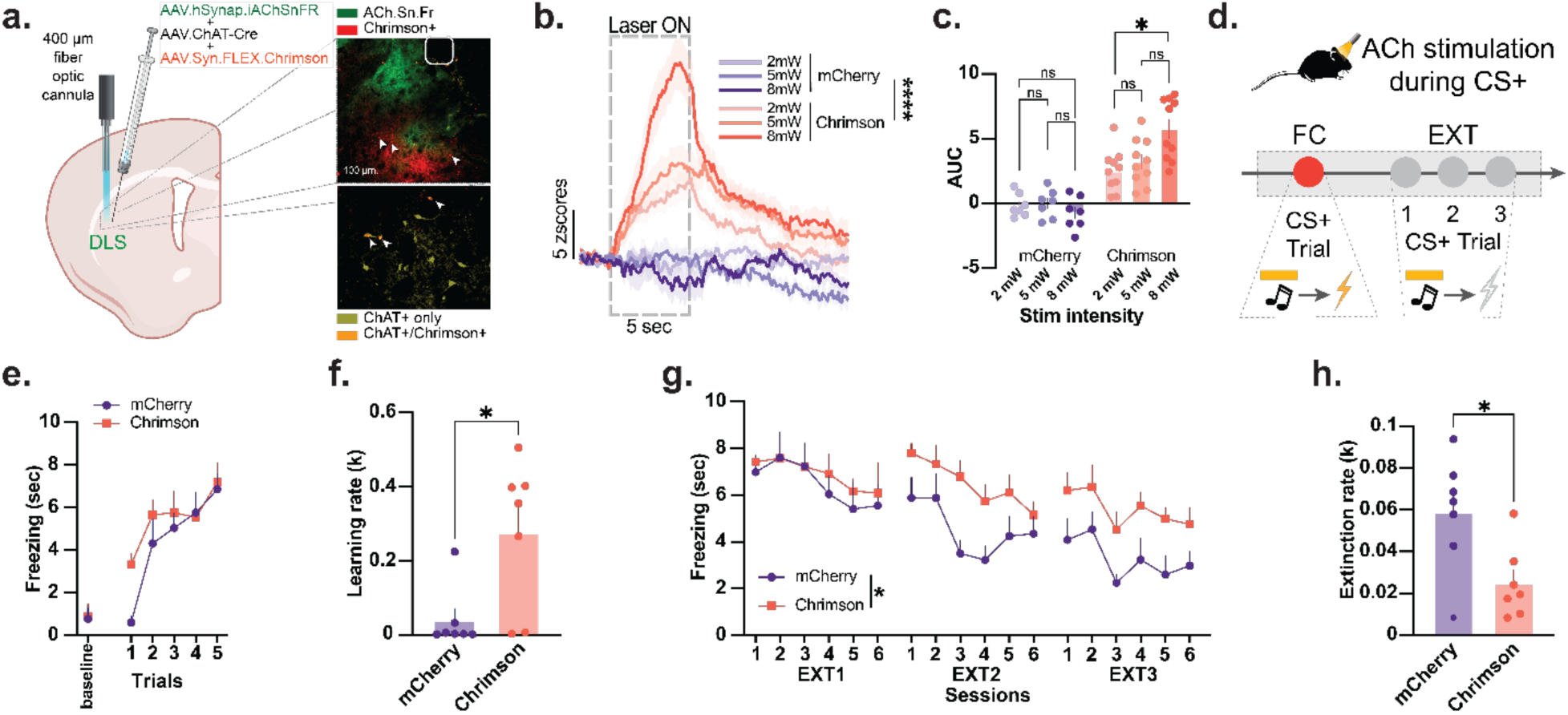
Optogenetic stimulation of dorsolateral striatal acetylcholine release enhances threat learning and impairs extinction. ***(a)*** Mice (n=14 mice, 7 males and 7 females) received unilateral DLS injections of a viral mixture containing AAV1.hSynap.iACh.Sn.FR, AAV2/5-ChAT-Cre-WPRE-HGH, and either AAV5-Syn-FLEX-rc[ChrimsonR-tdTomato] or AAV5-hSyn-DIO-mCherry. This strategy enabled optogenetic activation of ChAT+ DLS neurons (red, pointed with an arrow) while simultaneously monitoring ACh release with iACh.Sn.FR (green). Representative histology confirming co-expression is shown. ***(b)*** Optogenetic stimulation (589 nm) delivered at 2, 5, and 8 mW produced robust, dose-dependent increases in ACh release in Chrimson mice but not in mCherry controls. ***(c)*** Quantification revealed a significant increase in ACh signal in the *Chrimson* group mice (Two-Way RM ANOVA; Group main effect: F(1, 15)=70.94, *p*<0.0001; Sidak post-hocs: Chrimson 2mW vs 8mW, *p*=0.0178). ***(d)*** Mice then underwent one fear-conditioning session followed by three extinction sessions, with optogenetic stimulation applied during each CS+ presentation. ***(e–f)*** During conditioning, Chrimson animals acquired fear conditioning across trials and learned the CS–US association more quickly than mCherry controls (Learning rates: Unpaired t-test; *t*(12)=2.924, *p*=0.0127; n=7 mice per group). ***(g-h)*** During extinction, Chrimson animals exhibited persistently elevated freezing across sessions, indicating impaired extinction learning. Chrimson mice showed significantly reduced extinction rates, demonstrating that elevated cholinergic activity during threat cues biases learning toward sustained threat expectancy (Extinction rates: Unpaired t-test; t(12)=2.792, p=0.0163; n=7 mice per group). Data represented as mean ± S.E.M. ** p < 0.01.

First, before testing learning effects, we assessed whether optogenetically induced DLS ACh release was intrinsically aversive. In a conditioned place aversion (CPA) test (**Extended Data Fig. 6**), *Chrimson* and *mCherry* animals showed no differences in chamber preference or immobility (**Extended Data Fig. 6b,c**), indicating that DLS ACh stimulation did not induce place aversion. Similarly, open-field testing (**Extended Data Fig. 7**) revealed no group differences in total distance traveled or immobility (**Extended Data Fig. 7b,c**), demonstrating that stimulation did not alter locomotion.

We next evaluated whether optogenetically elevating DLS ACh during CS+ presentations affected fear learning and extinction (see **Fig. 3d** for a schematic). *Chrimson* animals displayed significantly enhanced acquisition of fear learning relative to *mCherry* controls (**Fig. 3e,f**), indicating that increasing DLS ACh release during CS+ accelerates threat learning. During extinction, *mCherry* animals showed the expected decline in freezing as CS+ no longer predicted foot-shocks, whereas *Chrimson* animals maintained high freezing levels (**Fig. 3g**) and showed slower extinction rates (**Fig. 3h**). These findings suggest that optogenetically elevated DLS ACh release during the CS+ impairs the updating of threat predictions. Overall, these results suggest that DLS ACh causally modulates threat perception during fear learning and extinction without inducing secondary aversive associations or locomotor deficits.

### ACh release alters DLS neural ensemble recruitment under baseline conditions

Our results demonstrate that ACh release in the DLS contributes to threat perception; however, how this threat-evoked cholinergic signaling reshapes DLS neuronal ensemble activity during threat learning remains unclear. To address this question, we employed dual-function nVoke miniature microscopes (Inscopix), which enable simultaneous optogenetic manipulation of DLS cholinergic interneurons (CINs) and single-cell calcium imaging of local neuronal ensembles in awake, behaving mice. This approach provides the unique ability to directly link cholinergic modulation to ensemble-level dynamics underlying threat learning, an experimental capability not achievable with traditional photometry or slice-based methods. To accomplish this, we injected a calcium sensor (AAV1.CaMK2a.GCaMP6m.WPRE.SV40) into the DLS to monitor the ensemble activity and used the same Cre-dependent optogenetic excitatory viral strategy used in experiments above (co-expression of ChAT-Cre and FLEX-Chrimson [*Chrimson* animals] or DIO-mCherry [*mCherry* animals]) to enable CIN excitation via optogenetic stimulation.

First, we examined how optogenetically evoked ACh release alters DLS neural ensemble calcium transients under baseline conditions. Initial calcium recordings confirmed reliable detection of CINs based on their robust, time-locked activity levels during optogenetic stimulation (10Hz, 5 mW; 10sec; **Fig. 4a**). In the *Chrimson* group, CINs constituted 1.91% of the imaged neuronal ensembles as they showed intense activity during the LED stimulation in a time-locked manner, and the rest consisted of other neuron types, named as non-cholinergic neurons (non-CINs; 98.09%; see **Extended Data Fig. 8** for the validation of our CIN identification method). We compared the activity of the CINs and non-CIN cells during the pre-LED period (10 seconds before the LED stimulation of CINs) and during the LED period. We observed a large-amplitude optogenetically evoked calcium response in CINs compared to non-CIN neurons during the LED period (**Fig. 4b-d**). We leveraged this high-threshold response *(∼10x of non-CIN cell calcium levels)* to identify CIN cells. In the *Chrimson* group, optogenetic stimulation increased the proportion of excited non-CIN cells (from 3.70% to 10.33%, **Fig. 4e**), whereas the fraction of inhibited and rebound-type cells (silent during LED stimulation with an immediate increase in activity at the LED offset) remained relatively stable. Hierarchical clustering analysis revealed a similar organization based on the LED-induced activity patterns in the non-CIN cells (**Fig. 4f,g**). Moreover, *mCherry* controls showed no detectable LED-induced changes in ensemble activity patterns (**Extended Data Fig. 9a,b,c**). Notably, optogenetic activation of CIN ACh release in Chrimson mice produced a clear increase in baseline calcium activity in non-CIN neurons compared to controls (**Extended Data Fig. 9d**).

**Figure 4.**
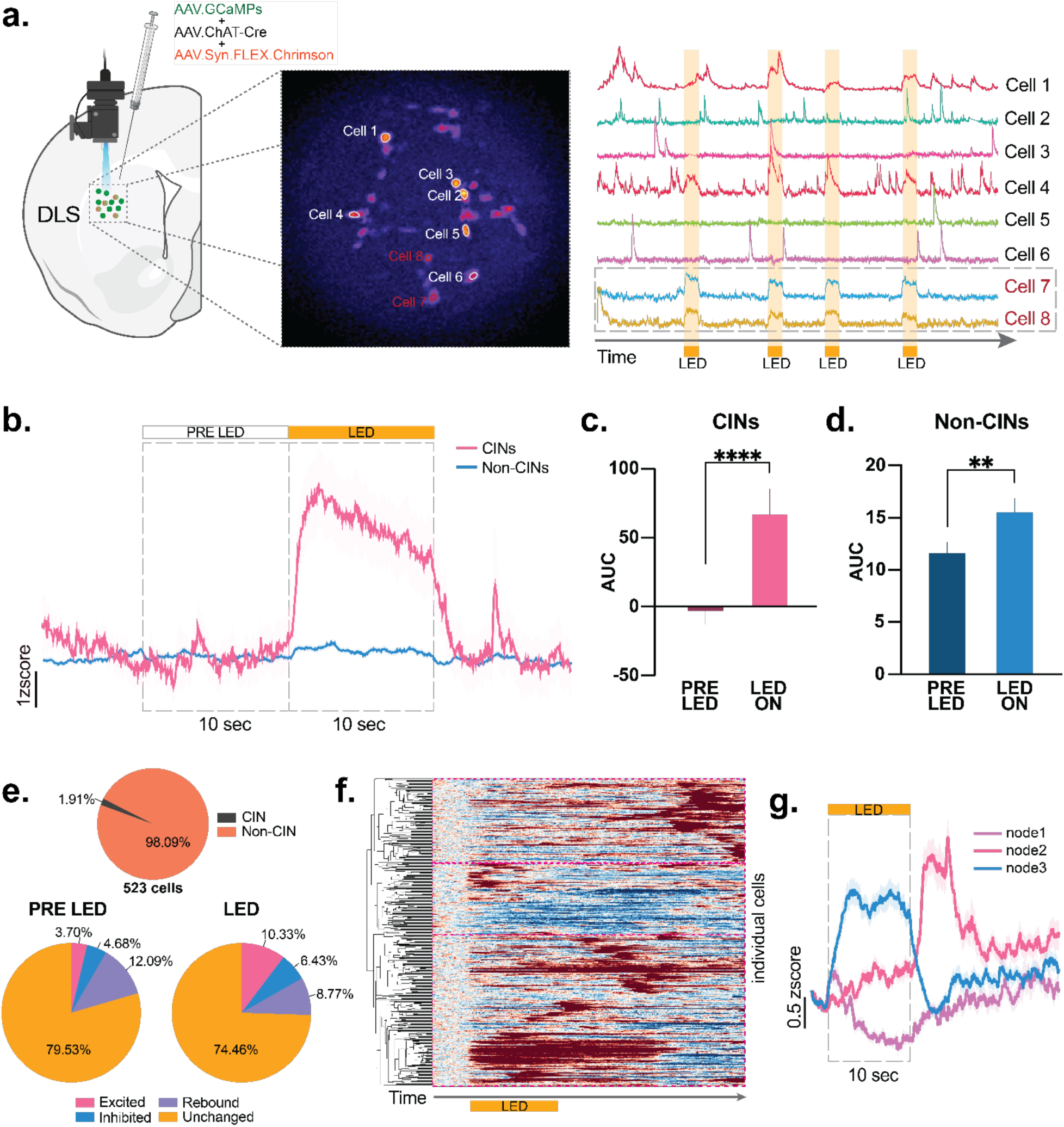
Single-cell calcium imaging reveals ensemble-level responses to optogenetic activation of DLS cholinergic interneurons. ***(a)*** Mice (n=14 mice, 7 males and 7 females) received unilateral DLS injections of AAV1.CaMKIIa.GCaMP6m.WPRE.SV40, AAV2/5-ChAT-Cre-WPRE-HGH, and either AAV5-Syn-FLEX-rc[ChrimsonR-tdTomato] or AAV5-hSyn-DIO-mCherry. This approach enabled simultaneous in vivo calcium imaging of DLS neurons and optogenetic activation of ChAT+ CINs. Representative fields of view illustrate identified CINs (red) and non-CIN (white/black) neurons responsive to optical stimulation. ***(b)*** Average traces from CINs and non-CINs during the pre-LED and LED periods in Chrimson mice show that optogenetic activation produces supra-physiological calcium activity in CINs. We leveraged this high-threshold response (∼10x of non-CIN cell calcium levels) to identify CIN cells. ***(c–d)*** Quantification of ΔF/F signals revealed robust increases in calcium activity during LED stimulation for both CINs (Paired t-test; *t*(39)=4.446, *p*<0.0001; n=10 cells x 4 LED stims = 40 LED stimulations) and non-CINs (Paired t-test; *t*(2051)=3.137, *p*=0.0017; n=513 cells x 4 LED stims = 2052 LED stimulations) compared to pre-LED baseline. ***(e)*** Across 523 imaged neurons, 1.91% were identified as CINs and 98.09% as non-CINs. Overall, DLS single-cells illustrate heterogeneous responses during LED stimulation, including cells activated by stimulation (Excited), cells inhibited during LED (Inhibited), and cells showing post-stimulation rebound dynamics. In Chrimson mice, the proportion of neurons excited during LED stimulation increased from 3.70% at baseline to 10.33% during stimulation. In mCherry controls, 5.16% showed excitation. ***(f)*** Hierarchical clustering of single-cell calcium signals during the LED stimulation period (Chrimson mice only) identified distinct response nodes. The heatmap shows all neurons ordered by cluster membership, revealing multiple subpopulations with robust LED-evoked activation following CIN stimulation. ***(g)*** Averaged ACh calcium transients from the top three response nodes (LED period, Chrimson mice only) show distinct activation motifs, with each node exhibiting a characteristic response profile during optogenetic CIN stimulation. These patterned increases indicate that CIN activation selectively recruits structured ensembles within the DLS. Data represented as mean ± S.E.M. ** p < 0.01, **** p < 0.0001.

Together, these findings indicate that time-locked increases in DLS ACh release, achieved via optogenetic stimulation of the DLS CINs under baseline behavioral conditions, enhance overall activity of the DLS neural ensembles.

### DLS ACh release primes local neural ensembles for upcoming threat

Next, we examined how optogenetically enhancing DLS ACh release shapes DLS neural ensembles dynamics during threat perception. First, we monitored single-cell DLS ensemble activity during a fear conditioning paradigm in which CINs were optogenetically stimulated during each CS+ presentation (see **Fig. 5a** for a schematic description). Consistent with our above results, *Chrimson* animals exhibited accelerated fear acquisition, exhibiting steeper fear learning curves compared to *mCherry* controls (**Fig. 5b,c**). CS+-evoked activity in DLS ensembles was comparable across *Chrimson* and *mCherry* groups (**Fig. 5d,e**). Interestingly, however, non-CIN cells in the *Chrimson* group displayed significantly reduced activity during the footshock compared to *mCherry* controls (**Fig. 5d,f**), suggesting that elevated DLS cholinergic activity reshapes ensemble organization in a manner that dampens footshock-evoked neural responses.

**Figure 5.**
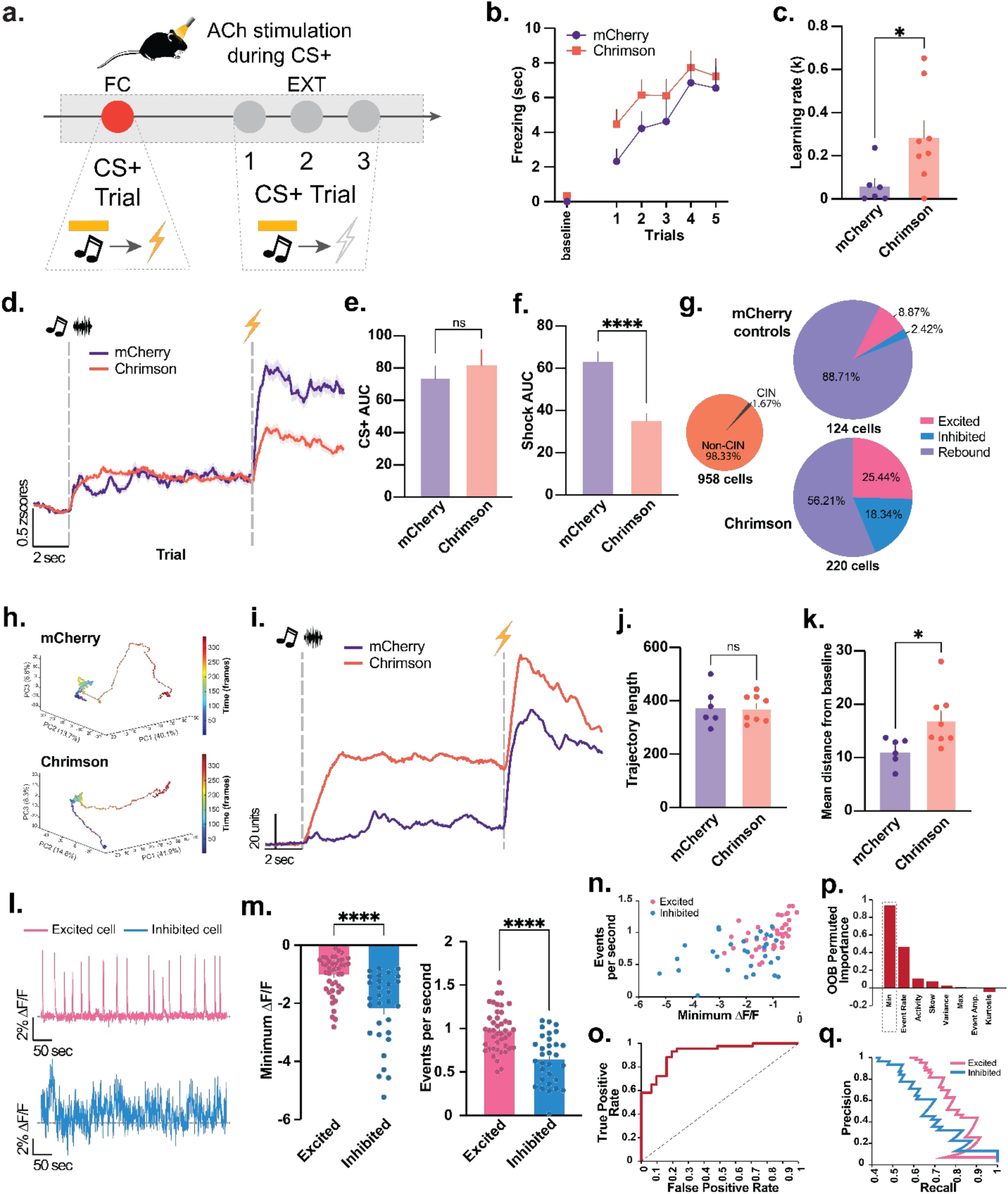
DLS ACh release primes local neural ensembles for the upcoming threat. ***(a)*** Mice (n=14 mice, 7 males and 7 females) underwent one fear-conditioning session followed by three extinction sessions. During conditioning, animals received six CS+ presentations (85 dB tone or white noise, counterbalanced; 10 s) paired with a footshock (0.5 mA, 0.5 s). During extinction, animals received twelve unreinforced CS+ presentations. DLS ChAT+ neurons were optogenetically stimulated during each CS+ presentation. ***(b–c)*** Chrimson mice exhibited elevated freezing during CS+ presentations and acquired the cue–shock association more rapidly than mCherry controls (Learning rates: Unpaired t-test; *t*(12)=2.338, *p*=0.0375; n=6-8 mice per group). ***(d)*** Population calcium activity aligned to CS+ and footshock revealed distinct neural response patterns between groups. ***(e)*** CS-evoked calcium activity did not differ significantly between Chrimson and mCherry control mice (Welch’s t-test; *t*(1388)=0.6560, *p*=0.5119; n= 462-958 cells per group). ***(f)*** Shock-evoked calcium responses were significantly reduced in Chrimson animals (Welch’s t-test; *t*(1024)=4.967, *p*<0.0001; n= 462-958 cells per group), indicating that CIN activation dampens the neural encoding of the aversive event itself. ***(g)*** Across 958 imaged DLS neurons, 1.67% were classified as CINs and 98.33% as non-CINs. In Chrimson mice, 25.44% of the Non-CINs that showed a CS+ response (excitation, inhibition, or rebound) were excited by optogenetic stimulation compared to 8.87% in mCherry controls. The proportion of inhibited cells was markedly higher during stimulated ACh release in Chrimson mice (18.34% vs. 2.42% in mCherry controls), indicating robust reorganization of DLS neural ensemble activity patterns. ***(h-i)*** State-space trajectory analysis of DLS ensemble activity during CS+ trials revealed divergent neural trajectories between groups. ***(j)*** The trajectory length did not differ significantly between mCherry and Chrimson mice (Welch’s t-test; *t*(8.614)=0.1547, *p*=0.8806; n = 6-8 mice per group). ***(k)*** The mean distance from baseline was significantly greater in Chrimson animals (Welch’s t-test; *t*(10.63)=2.699, *p*=0.0213; n = 6-8 mice per group), indicating that optogenetic CIN activation reorganizes DLS ensembles by redistributing neurons into strongly excited and inhibited subpopulations rather than simply scaling overall activity. ***(l)*** Representative single-cell calcium traces (LED period removed for clarity) from neurons identified as excited or inhibited during optogenetically stimulated ACh release. ***(m)*** Inhibited neurons exhibited significantly lower minimum ΔF/F values (Welch’s t-test; *t*(40)=4.571, *p*<0.0001; n= 30-43 cells per group; One cell was excluded as an outlier) and produced fewer calcium events than excited neurons (Welch’s t-test; *t*(59.02)=5.119, *p*<0.0001; n= 31-43 cells per group), indicating distinct physiological response profiles. ***(n)*** Pairwise scatter plots of event rate and minimum ΔF/F illustrate clear separation between excited and inhibited neurons in feature space. ***(o)*** ROC curves for generalized linear models (GLMs) trained on baseline trace features show strong discriminability between excited and inhibited cells (χ² improvement over constant model = 48.2; *p*<0.0001; GLM LED-free baseline accuracy = 86.49%). ***(p)*** Random forest feature-importance analysis revealed that minimum ΔF/F was the most informative predictor (OOB permuted importance = 0.93), followed by event rate (OOB permuted importance = 0.57). ***(q)*** Precision–recall curves for the random forest classifier demonstrate high classification performance for both excited and inhibited neuronal populations. Data represented as mean ± S.E.M. * p < 0.05, **** p < 0.0001.

Although the overall CS+-evoked responses of DLS neurons did not differ between groups, the underlying ensemble organization was markedly more dynamic in *Chrimson* animals, indicating enhanced network reorganization despite comparable mean activity levels. Specifically, the *Chrimson* group ensembles showed distinct activity patterns following CS+ onset compared to the *mCherry* group, an increased number of both excited (*mCherry*: 8.87% *Chrimson*: 25.44%) and inhibited (*mCherry*: 2.42% *Chrimson*: 18.34%) neurons, and a decreased number of rebounded neurons (*mCherry*: 88.71% *Chrimson*: 56.21%; **Fig. 5g**; see **Extended Data Fig. 10** for averaged cell traces for each response group). State-space trajectory analyses further revealed that optogenetic enhancement of ACh release during CS+ presentations increased the divergence of DLS ensemble activity from baseline, reflecting a reorganization of population dynamics rather than a uniform elevation in single-cell firing (**Fig. 5h-k**).

Finally, to determine whether the polarity of non-CIN responses to optogenetic CIN stimulation was already present in their spontaneous activity, we analyzed LED-free baseline traces from all recorded non-CIN neurons. The LED periods-removed baseline traces of excited and inhibited cells in the *Chrimson* group showed clearly distinct activity patterns (**Fig. 5l**; **Extended Data Fig. 11a**). Neurons that were ultimately inhibited displayed deeper negative baseline minima and lower spontaneous event rates than those that later showed excitation during optogenetic ACh release (**Fig. 5m**). Importantly, the two groups did not differ in their baseline or peak activity levels, indicating that the neurons classified as “inhibited” were not inherently less active at baseline compared to the neurons that were later excited (**Extended Data Fig. 11b,c**). Excited and inhibited cells formed clearly separable clusters in baseline feature space, as shown by pairwise scatter plots (**Fig. 5n**). A generalized linear model trained solely on baseline features accurately predicted which neurons would become excited or inhibited (86.5% accuracy; χ² = 48.2), demonstrating that spontaneous activity patterns contained strong predictive information (**Fig. 5o**). LASSO-regularized regression and Random Forest permutation-importance analyses independently confirmed that baseline activity could reliably predict whether a neuron would become excited or inhibited. Both models converged on the same dominant predictors: the baseline minimum was by far the strongest determinant of response polarity (importance = 0.93), followed by spontaneous event rate (0.47), with baseline mean, skewness, and variance contributing more modestly (**Fig. 5p**). Consistent with these feature-importance profiles, the Random Forest classifier also successfully predicted response polarity, as reflected in its precision–recall performance (**Fig. 5q**), indicating that CIN-evoked re-organization of DLS neural ensembles leverages the pre-existing functional states of non-CIN neurons. Cells with deeper negative excursions and elevated spontaneous event rates are biased toward inhibition, whereas neurons that maintain higher tonic levels and more stable baseline dynamics are biased toward excitation during CIN activation.

Together, these results indicate that optogenetically enhanced ACh release primes DLS neural ensembles for the upcoming threat by dynamically reorganizing their activity patterns during the predictive CS+ period in a manner that reflects and capitalizes on the neurons’ intrinsic spontaneous activity profiles of the local circuitry.

### Elevated DLS ACh release maintains threat expectancy and impairs fear extinction

Finally, we investigated how optogenetically elevated ACh release modulates DLS neural ensemble dynamics during fear extinction, leading to the delayed extinction of fear memories. Replicating our initial optogenetic results, *Chrimson* animals again exhibited impaired extinction learning (**Fig. 6a, b**). During the final extinction session (EXT3), optogenetically elevating ACh release at CS+ onset profoundly reorganized DLS ensemble activity patterns. *Chrimson* animals maintained a markedly elevated proportion of excited neurons (24.14%) compared to *mCherry* controls (1.79%), along with a modestly higher proportion of inhibited neurons (26.72% vs. 19.64%; **Fig. 6c**). Thus, even at the end of extinction, enhanced cholinergic signaling sustains a broad and polarized ensemble response, preventing the DLS network from returning to the subdued activation state observed in controls.

**Figure 6.**
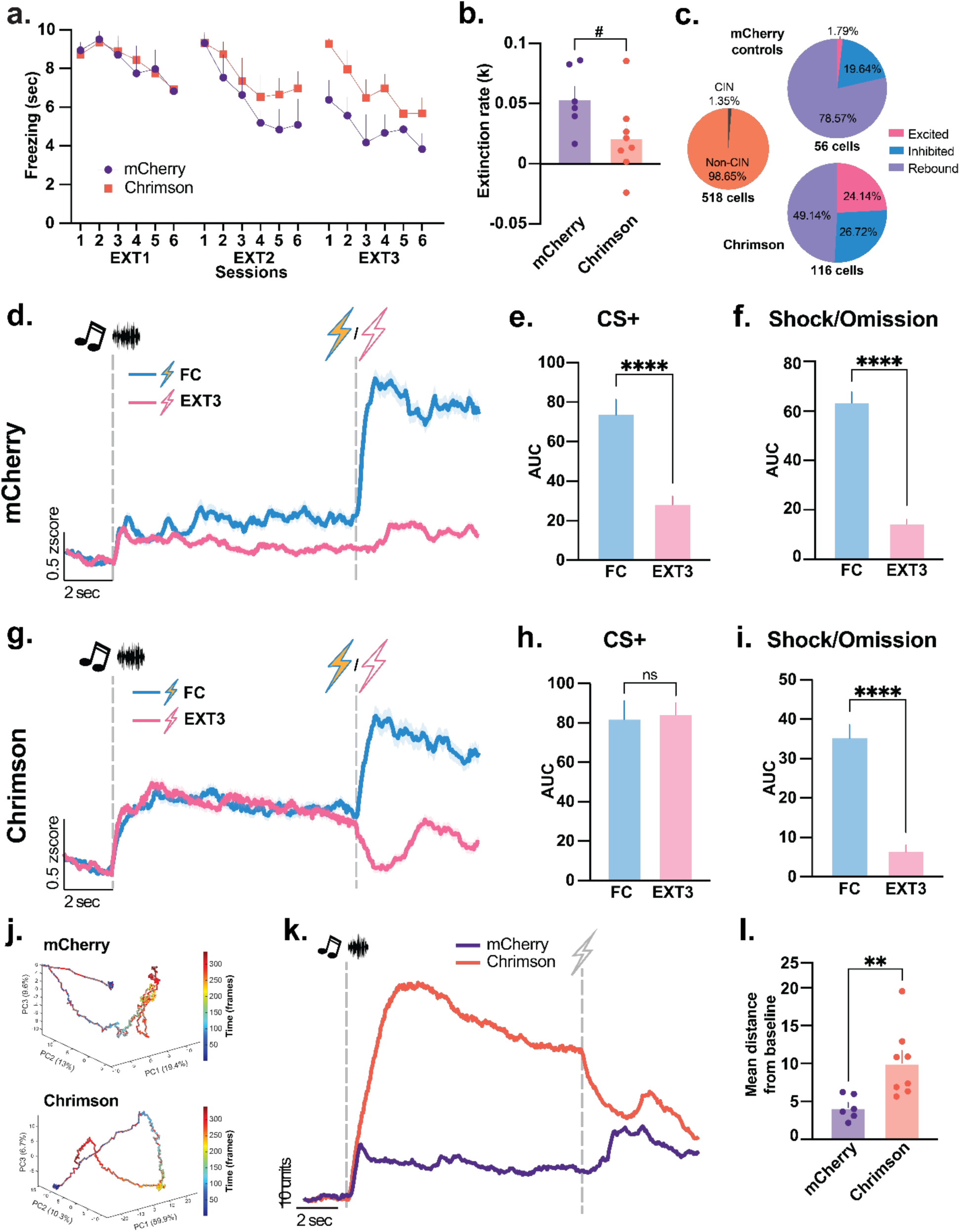
Elevated DLS ACh release maintains threat expectancy and impairs fear extinction. ***(a-b)*** Across extinction sessions (Ext1, Ext2, Ext3), Chrimson mice maintained higher freezing levels than mCherry controls, indicating delayed extinction. Learning indices confirmed that Chrimson animals were significantly slower to learn that the CS+ no longer predicted footshock (Extinction rates: Unpaired t-test; *t*(12)=2.012, *p*=0.0672; n=6-8 mice per group). ***(c)*** During extinction, a substantially greater proportion of the Non-CINs that showed a CS+ response (excitation, inhibition, or rebound) were excited by optogenetic stimulation in Chrimson mice (24.14%) compared to mCherry controls (1.79%), reflecting altered ensemble recruitment. ***(d)*** In mCherry controls, DLS calcium activity decreased across learning. ***(e)*** CS+-evoked calcium responses were significantly lower in extinction session 3 than during fear conditioning (Welch’s t-test; *t*(789.6)=4.808, *p*<0.0001; n= 394-462 cells per group). ***(f)*** Likewise, shock omission responses during extinction session 3 were significantly reduced relative to shock responses during conditioning (Welch’s t-test; *t*(673.7)=9.819, *p*<0.0001; n= 394-462 cells per group). ***(g)*** In Chrimson mice, calcium traces aligned to CS+ onset and shock/omission revealed distinct patterns between conditioning and extinction. ***(h)*** CS+-evoked calcium activity did not differ significantly between fear conditioning and extinction session 3 (Welch’s t-test; *t*(907.8)=0.1227, *p*=0.9024; n= 518-958 cells per group). ***(i)*** However, shock omission evoked a markedly reduced calcium response during extinction session 3 compared to the robust shock response during conditioning (Welch’s t-test; *t*(1461)=6.377, *p*<0.0001; n= 518-958 cells per group). ***(j–k)*** State-space trajectory analysis of DLS calcium dynamics in Chrimson and mCherry mice during extinction session 3 showed distinct ensemble trajectories between groups, indicating altered population structure with CIN activation. ***(l–m)*** Distance-from-baseline analysis demonstrated that Chrimson mice displayed significantly greater deviations from baseline ensemble activity during extinction session 3 (Welch’s t-test; *t*(9.103)=3.329, *p*=0.0087; n = 6-8 mice per group), reflecting CIN-driven reorganization of DLS network states during omission. Data represented as mean ± S.E.M. # p = 0.0672, ** p < 0.01, **** p < 0.0001, ns= not significant.

This pattern was also apparent in the evolution of mean calcium responses in DLS neuronal ensembles from fear conditioning to the final extinction session (EXT3), which revealed a striking divergence between groups. In *mCherry* controls, ensemble activity to both the CS+ (**Fig. 6d,e**) and the omitted footshock progressively decreased as extinction proceeded (**Fig. 6d,f**). In sharp contrast, this adaptive reduction was absent in the *Chrimson* group. Despite the absence of an actual threat, optogenetically elevated ACh release in the DLS elicited persistent ensemble activation to the previously threat-predictive cue (CS+, **Fig. 6g,h**), effectively maintaining a heightened threat-expectancy state. Moreover, when the anticipated footshock did not occur, DLS ensemble activity exhibited a marked decrease (**Fig. 6g,i**), resembling a negative-prediction error-like response. Across both early (EXT1) and late (EXT3) extinction session, non-CIN neurons in *Chrimson* animals exhibited a stronger response to the CS+ and a weaker response to the omitted footshock compared to controls (**Extended Data Fig. 12**). These findings indicate that optogenetically enhanced ACh release biases DLS non-CIN ensembles toward maintaining threat-predictive activity even in the absence of actual threat, thereby sustaining an expectancy of aversive outcomes during extinction.

State-space trajectory analyses further revealed that elevated ACh release in the DLS produced a significantly greater divergence of ensemble activity from baseline (**Fig. 6j–l**), signifying a robust, sustained reorganization of population dynamics that reinforces threat expectancy and hinders extinction. Together, these findings demonstrate that heightened DLS ACh tone reshapes striatal network states to favor persistent threat encoding, reducing behavioral flexibility and impairing the suppression of previously learned fear.

## Discussion

Here, we established that ACh release in the DLS functions as a valence-biased signal that preferentially signals aversive experiences. Across behavioral paradigms, ACh increased during threat and decreased during reward prediction, revealing a robust negative-valence tuning of DLS cholinergic activity. DLS ACh release was causal for threat memory encoding, as optogenetic excitation of DLS CINs enhanced fear learning and impaired fear extinction without producing aversion itself. Concurrent single-cell imaging revealed that CIN activation reorganized DLS ensemble dynamics, sustaining threat expectancy during both conditioning and extinction. Unlike controls, which displayed strong shock-evoked responses and rapidly extinguished both neural and behavioral reactions, stimulated ACh release dampened shock responses while maintaining cue-evoked activation, indicating that cholinergic signaling primes DLS ensembles for impending threat. Importantly, this cholinergic modulation was not uniform. Baseline activity profiles predicted whether individual non-CIN neurons would be excited or inhibited by CIN activation, demonstrating that ACh recruits pre-existing functional states within the DLS network. Overall, these findings establish DLS ACh release as a key modulator of striatal network dynamics that biases learning toward threat persistence.

Our results establish DLS ACh release as a mechanism that relocates cognitive resources toward threat perception. Indeed, ACh has long been recognized as a neuromodulator of attention and learning, regulating cortical and hippocampal network states to optimize information processing^37–40^. Within the striatum, the primary source of ACh is local CINs, which interact with dopaminergic and GABAergic systems to influence reinforcement learning and action selection^19–21^. Here, we define the affective dimension of striatal ACh signaling, revealing a selective role for DLS ACh in encoding negative valence. Our results demonstrate that ACh release increases during aversive experiences and decreases during reward consumption, providing a bidirectional, valence-specific signal that shapes behavioral outcomes. This work extends prior evidence implicating that increased cholinergic input from the basal forebrain to regions such as the hippocampus, amygdala, and prefrontal cortex contributes to negative affective states, including anxiety-, depression-like, and threat-related behaviors^41–45^. However, our findings contrast with ACh release patterns in cortical regions. For example, stimulating basal forebrain cholinergic terminals in the medial prefrontal cortex (mPFC) during CS+ presentations does not alter aversive CS–footshock learning^46^. This suggests that striatal CIN-derived local ACh represents a region-specific, functionally distinct cholinergic pathway tuned to signal valence and guide learning in ways not shared by basal forebrain projections.

We also identify the DLS as a critical subcortical locus of valence-dependent signaling, mediated by local CIN-derived ACh, which dynamically shapes striatal neural ensemble organization. Traditionally, the DLS has been viewed primarily as a motor and habit-related structure engaged during overtrained, stimulus–response learning^27,29,47^. Here, we demonstrate that DLS neural ensembles not only respond to aversive stimuli but also reorganize in anticipation of threat. This anticipatory function likely reflects an ACh-driven reorganization of local circuit dynamics, in which heightened cholinergic signaling shifts ensemble activity toward threat-predictive states, enhancing neurons that encode danger while dampening those that signal safety or reward. Notably, our results show that optogenetic CIN stimulation did not simply uniformly amplify overall DLS responses to the CS+ during fear conditioning. Instead, elevated ACh increased the proportion of both excited and inhibited neurons, indicating a reorganization of ensemble composition within the DLS, rather than a simple gain change. These findings suggest that ACh release during threat exposure restructures DLS circuitry, tuning it toward heightened threat sensitivity and anticipation.

Given the dense and broadly distributed nature of cholinergic signaling within the striatum^17,48^, multiple plausible pathways likely contribute. It is well established that CIN-derived ACh orchestrates local ensemble dynamics through differential activation of nAChRs and mAChRs. Specifically, mAChRs-mediated signaling may dampen excitatory inputs to medium spiny neurons (MSNs)^49–52^ or modulate inhibitory transients through GABAergic interneurons^53–56^. In contrast, the activation of nAChRs on dopaminergic and glutamatergic terminals and GABAergic interneurons may enhance the release of dopamine^57–59^, glutamate^60,61^, and GABA^55,62^ onto MSNs, respectively. Our and others’ prior work have demonstrated that nAChR signaling is critical for aversive learning: systemic nicotine, an nAChR agonist, enhances contextual and trace fear conditioning^63–65^, delays extinction of contextual fear^66–69^, and enhances spontaneous recovery of extinguished fear^70^ through the high-affinity α4β2 nAChRs. Through these receptor-specific, bidirectional signaling pathways, CIN-derived ACh is positioned to dynamically tune excitatory and inhibitory balance and ensemble synchrony that govern valence-dependent encoding, biasing striatal networks towards aversive states.

In line with this framework, our data reveal that the polarity of non-CIN responses likely reflects underlying cell-type–specific mechanisms within the DLS. Although we did not genetically identify the recorded neurons, the observed response profiles align with known cholinergic effects in this region. Neurons excited by CIN stimulation likely correspond to MSNs depolarized by M1 muscarinic receptor–mediated facilitation of intrinsic excitability^71^. In contrast, the inhibited population may include GABAergic interneurons, such as PV⁺ fast-spiking or NPY⁺ low-threshold spiking cells, which are hyperpolarized through M2/M4 muscarinic signaling^72–74^, and a subset of inhibited neurons could also represent D1-MSNs expressing M4 receptors, which oppose D1-dependent cAMP signaling and suppress excitability^75,76^. Thus, CIN-evoked reorganization of DLS ensembles likely exploits these receptor-specific mechanisms to transiently bias the local network toward MSN excitation and interneuron suppression, effectively gating striatal output toward threat-related encoding.

By linking cholinergic activity to both emotional valence and circuit- and population-level reconfiguration, our findings position DLS ACh as a fundamental component of the brain’s threat-processing architecture. Rather than merely facilitating learning or attention, striatal cholinergic signaling appears to define an internal state of vigilance, biasing the organism toward avoidance and anticipatory threat evaluation. Dysregulated DLS Ach signaling may therefore contribute to maladaptive threat learning and persistent avoidance behaviors characteristic of anxiety and stress-related disorders. By identifying a striatal cholinergic mechanism that integrates valence encoding with behavioral flexibility, our findings highlight dorsolateral striatal ACh as a potential therapeutic target for restoring adaptive affective processing.

## Acknowledgements

This work was supported by NIH grant MH132052 to MGK and T32DA007237 to BPP and the Brain and Behavior Research Foundation Young Investigator Award to MGK.

## Author contributions

MGK and OD designed the behavioral and imaging studies. OD, NHW, CM, TG, AC, BPP, and FZB executed the single-cell imaging, fiber photometry, and optogenetic experiments, including surgeries, behavioral and imaging experiments, and histology. MGK, OD, and NHW analyzed the data. MGK, OD, and NHW wrote the manuscript. All authors edited and reviewed the manuscript.

## Competing Interests

The authors have nothing to disclose.

**Correspondence and requests for materials should be addressed to Munir Gunes Kutlu**.

**Supplementary Information is available for this paper.**

## Methods

### Data availability

Data reported in this paper are available from the lead contact upon reasonable request.

### Code availability

The custom codes used for analyzing the datasets reported in this study will be provided upon reasonable request.

### Animals

Male (n=29) and female (n=22) 8-week-old C57BL/6J mice obtained from Jackson Laboratories (Bar Harbor, ME; Strain #:000664) were kept 2-4 per cage and maintained on a 12-hour reverse light/dark cycle, with all behavioral testing taking place during the dark cycle. Animals were given ad libitum access to food and water. All experiments were conducted in accordance with the guidelines of the Institutional Animal Care and Use Committee (IACUC) at Rowan University Virtua School of Osteopathic Medicine and Temple University Lewis Katz School of Medicine.

### Surgical Procedures

At least 1 hour prior to surgery, mice were administered Meloxicam (2 mg/kg) via subcutaneous injection. Animals were anesthetized using isoflurane (5% for induction and 2% for maintenance) and placed on a stereotaxic frame (David Kopf Instruments). Ophthalmic ointment was continuously applied to the eyes throughout the surgical procedures. A midline incision was then made down the scalp, and a craniotomy was performed with a dental drill using aseptic technique. A 10-mL Nanofil Hamilton syringe (WPI) with a 34-gauge beveled metal needle was used to infuse viral constructs into the dorsolateral striatum (DLS).

#### Fiber photometry and optogenetic surgeries

A fluorescent acetylcholine (ACh) sensor, pAAV.hSynap.iACh.Sn.FR (Addgene, #137950-AAV1), was unilaterally infused into the dorsolateral striatum (DLS; coordinates relative to the bregma: anterior/posterior, +0.38 mm; medial/lateral, −2.5 mm; dorsal/ventral, −3.3 mm) at a rate of 50 nL/min for a total volume of 500 nL. Following infusion, the needle was kept at the injection site for 7 minutes before being slowly withdrawn. In optogenetic surgeries, an even ratio mixture of rAAV-ChAT-Cre-WPRE-HGH polyAAV2/5 (Biohippo, #BHV12400208-3) and pAAV-Syn-FLEX-rc[ChrimsonR-tdTomato] (Addgene, #62723-AAV5) or pAAV-hSyn-DIO-mCherry (Addgene, #50459-AAV5) was infused into the DLS at a rate of 50 nL/min for a total volume of 700 nL following the ACh sensor infusion. Fiber-optic cannulas (400 μm core diameter; 0.48 NA; Doric) were then implanted in the DLS and positioned immediately dorsal to the viral injection site before being permanently fixed to the skull using adhesive cement (C&B Metabond; Parkell). **This method allowed us to verify that optogenetic stimulation of ChAT interneurons consistently evoked a reproducible, time-locked increase in acetylcholine release across animals.** Follow-up care was performed according to IACUC. Animals were allowed a minimum of 5 weeks to recover to ensure efficient viral expression before commencing experiments.

#### In vivo calcium imaging via miniscopes

A fluorescent calcium sensor, AAV1.CaMK2a.GCaMP6m.WPRE.SV40 (Inscopix, #1000-002977) was unilaterally infused into the DLS (coordinates relative to the bregma: anterior/posterior, +0.38 mm; medial/lateral, −2.5 mm; dorsal/ventral, −3.3 mm) at a rate of 50 nL/min for a total volume of 500 nL. Following infusion, the needle was kept at the injection site for 7 minutes before being slowly withdrawn. In optogenetic surgeries, an even ratio mixture of rAAV-ChAT-Cre-WPRE-HGH polyAAV2/5 (Biohippo, #BHV12400208-3) and pAAV-Syn-FLEX-rc[ChrimsonR-tdTomato] (Addgene, #62723-AAV5) or pAAV-hSyn-DIO-mCherry (Addgene, #50459-AAV5) was infused into the DLS at a rate of 50 nL/min for a total volume of 700 nL following the calcium sensor infusion. Following the viral infusions, a 23-gauge needle with blunted tip was used to clear the track before the GRIN lens (0.5 mm diameter, 4.0 mm length, 0.5 NA; #1050-004417, Inscopix) implantation in the DLS, immediately dorsal to the viral injection site, before being permanently fixed to the skull using adhesive cement (C&B Metabond; Parkell). Follow-up care was performed according to IACUC. Animals were allowed a minimum of 5 weeks to recover to ensure efficient viral expression before commencing experiments.

### Apparatus

Fiber photometry, optogenetic, and *in vivo* calcium imaging recordings were carried out in related behavioral apparatuses, including operant conditioning boxes (Med Associates Inc., St. Albans, Vermont) and an open field (40×40×22 cm, Ugo Basile SRL, Gemonio, Italy). Each operant conditioning box was fitted with visual and auditory stimuli, including a standard house light, a white-noise generator, and a 16-tone generator capable of outputting frequencies between 1 and 20 kHz (85 dB).

### Histology

At the end of each of the fiber photometry, optogenetics, and single cell calcium imaging experiments outlined below, subjects were deeply anesthetized with an intraperitoneal injection of Ketamine/Xylazine (100mg/kg/10mg/kg) and transcardially perfused with 10 mL of PBS solution followed by 10 mL of cold 4% PFA in 1x PBS. Animals were quickly decapitated, the brain was extracted and placed in 4% PFA solution, and stored at 4 °C for at least 48 hours. Brains were then transferred to a 30% sucrose solution in 1x PBS and allowed to sit until brains sank to the bottom of the conical tube at 4 °C. After sinking, brains were sectioned at 40μm on a freezing sliding cryostat (Leica CM3050 S). Sections were stored in a cryoprotectant solution (7.5% sucrose + 15% ethylene glycol in 0.1 M PB) at −20 °C until immunohistochemical processing.

#### Validation of fluorescent sensors of ACh and calcium

Viral expressions were validated via immunohistochemical staining. We immunohistochemically stained all DLS slices with an anti-GFP antibody (chicken anti-GFP; Sigma, #06-896; 1:500 in 5% bovine serum albumin (BSA); room temperature overnight) to validate iACh.Sn.FR and GCaMP6m expression in the DLS. Sections were then incubated with secondary antibodies (GFP: rabbit anti-chicken CF 488A [Sigma, #SAB4600052]; 1:1000 in 5% BSA) overnight at 4 °C. After washing, sections were incubated for 5 min with DAPI (NucBlue, Invitrogen) to achieve counterstaining of nuclei before mounting in Prolong Gold/Fluoro-gel (Invitrogen). Fluorescent images were taken using a Keyence BZ-X800 inverted fluorescence microscope (Keyence), under a dry 10x objective (Nikon). The injection site location and the fiber implant or GRIN lens placements were determined via serial imaging in all animals. We identified sections that displayed the DLS, viral expression, and fiber optic tip.

#### Validation of the ChAT-Cre/FLEX-Chrimson dual viral approach to target ChAT+ interneurons

For the validation of the ChAT interneuron specificity of the ChAT-Cre viral approach we employed for our optogenetic studies, we injected an even ratio mixture of rAAV-ChAT-Cre-WPRE-HGH polyAAV2/5 (Biohippo, #BHV12400208-3) and pAAV-Syn-FLEX-rc[ChrimsonR-tdTomato] (Addgene, #62723-AAV5) was infused into the basal forebrain (BF; coordinates relative to the bregma: anterior/posterior, −0.34 mm; medial/lateral, −1.5 mm; dorsal/ventral, −5.3 mm) where the majority of the cells are ChAT+^1,2^. The viral mixture was infused at a rate of 50 nL/min for a total volume of 500 nL following the ACh sensor infusion. After the perfusion and preparation of the BF slices, we stained the Chrimson and ChAT+ positive cells using an anti-mCherry antibody (Goat anti-mCherry, Thermo Fisher #PA5-143590; 1:1000 in 5% BSA) for ChrimsonR-tdTomato and an anti-ChAT antibody (rabbit anti-ChAT, Thermo Fisher #PA529653; 1:1000 in 5% BSA) for ChAT. Sections were then incubated with secondary antibodies (for mCherry ab: donkey anti-goat CF 594, Sigma #SAB4600095; for ChAT ab: chicken anti-rabbit Alexa Fluor 488, Thermo Fisher # A-21441; 1:1000 in 5% BSA) overnight at 4 °C. The imaging was executed as explained above. We quantified the total number of ChAT+ neurons and calculated the percentage of ChrimsonR-expressing cells that overlapped with ChAT+ labeling using FIJI (ImageJ).

### Behavioral Methods

All behavioral experiments were recorded using a USB camera from a top-down angle (Stoelting, #60516; TDT iVn, iV2; Inscopix, nVision).

#### Non-contingent stimulus delivery

Animals were subjected to different non-contingent stimulus deliveries on different days to record the ACh release change for 30 minutes. Subjected cues were as follows: foot shock (1 mA, 0.5 sec, single delivery), free access to sucrose (30%), and quinine (0.32g/L) via a sipper.

#### Appetitive conditioning

Animals were tested for seven consecutive days and subjected to a conditioned stimulus (CS+, 85dB white noise or tone, counterbalanced, 10 sec in duration) which signals sucrose access for 10 sec. Each session consisted of 15 CS+ in pseudo-random order with randomized intervals (20sec, 30sec, 45sec, 50sec). To quantify conditioned approach behavior, we calculated the mouse’s distance to the sipper zone on a frame-by-frame basis from the DeepLabCut tracking data (see below). The mouse’s position was estimated as the mean x–y coordinates of two reliably tracked body points, and the distance to a user-defined polygon corresponding to the sipper zone was computed for each frame. The resulting distance trace was then segmented into Pre-CS+ (baseline period prior to cue onset) and CS+ (cue presentation) epochs for each trial. Mean distance during each epoch was calculated to obtain a Pre-CS+ distance and a CS+ distance. To assess learning, we computed an Approach Ratio defined as:

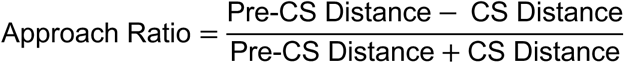

Higher Approach Ratio values reflect greater cue-evoked approach toward the sipper, indicating that the CS+ increasingly predicted sucrose reward.

#### Fear Conditioning/Extinction

Animals were tested consecutively in one fear conditioning session, followed by five extinction sessions. During fear conditioning, animals were subjected to CS+ (85dB white noise or tone, counterbalanced, 10 sec in duration), which signals foot shock (0.5 mA, 0.5 sec). The fear conditioning session consisted of 6 CS+ presentations in alternating order with randomized intervals (20sec, 30sec, 45sec, 50sec). During extinction sessions, animals were exposed to the same CS+ as in the fear conditioning session; however, foot shocks were omitted entirely. Freezing response was measured during both training and test sessions. The freezing response was defined as the time (seconds) that mice were immobile (lack of any movement, including sniffing) during the tone period and calculated as a percentage of the total cue time. We converted freezing durations to percentages of total cue time ((freezing time * 100)/ stimulus duration).

#### Positive Reinforcement and Punishment

Prior to positive reinforcement training, animals were exposed to 30% food restriction for the first three days, followed by 50% food restriction for the training and testing days. Body weights and food intakes were followed daily. At the end of the food restriction period, the animals’ diet returned to 100% ad libitum free access. Following 3 days of food deprivation, animals began their training to acquire 30% sucrose access via a sipper in operant boxes for 30 minutes. Once the 400 lick threshold was achieved, animals began training in a cued fixed-ratio 1 (FR1) protocol, in which each nose-poke response on the left nose-poke port within the cue (85 dB tone, 30sec, 10 trials per session) allowed 30% sucrose access (10sec). After animals reached 50% correct nose poke efficiency, animals started to receive a foot shock (0.3mA, 0.5sec) for each correct nose poke as well as 30% sucrose access (10sec).

#### Open Field Test

Animals were placed in the arena by themselves to move freely for two consecutive 5-minute trials. The average distance traveled and immobility were determined.

#### Condition Place Aversion (CPA)

The biased conditioned place preference/aversion paradigm protocol was adapted from earlier publications from our group^3^. The CPP training and testing were carried out in an apparatus comprising two distinct compartments with distinct sensory cues (smooth floor or floor with holes) to determine whether the optogenetic stimulation is aversive. Animals were allowed to explore freely both compartments alone for 10 min. Time spent in each compartment was recorded, and the preferred compartment was determined. The following day, the animals once again freely explored the apparatus, receiving optogenetic stimulation while they were in their preferred compartment. The change in time spent in each compartment was used to determine whether a triggered aversion was present or absent.

#### Deep Lab Cut markerless tracking analysis

We recorded each mouse’s movement using a USB camera (Stoelting, #60516; TDT iVn, iV2; Inscopix, nVision, 15-30 frames per second) attached above the operant box. We used DeepLabCut (Python 3, DLC, version 2.2b8, 112) for markerless tracking of position. DLC was trained on 250+ frames from multiple videos, and these frames were annotated and used to train a ResNet-50 neural network for 200000 iterations. The locations of the mice were computed as x/y coordinates converted into centroids, calculated as the average of x and y coordinates. We calculated the number of frames each mouse spent in:

1. The stimulated vs non-stimulated compartment for the CPA experiment
2. Around the sipper port (200px x 164px zone) for the appetitive conditioning experiments

For the open field experiment, we calculated the total distance moved and immobility. For each body part tracked by DeepLabCut, the script first extracted the x–y coordinates only when the tracking likelihood was ≥ 0.9. It then computed the frame-to-frame Euclidean displacement (i.e., the change in position between consecutive frames) for each body part:

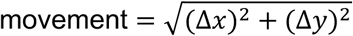

This produced a movement trace for each body part across the session. The script then summed the movement across all frames and averaged across body parts, giving the total distance traveled (in pixels). The instantaneous movement traces from all body parts were averaged to yield a single movement signal across the body. This signal was then smoothed using a 2-frame moving window to reduce noise. Frames where the smoothed movement value fell below a threshold of 2 pixels/frame were classified as immobile. Immobility was reported as the percentage of frames below this movement threshold.

#### Statistical analysis of behavioral measures in fiber photometry experiments

For fear conditioning, we analyzed freezing response to the CS+ between Trial 1 and Trial 6 using a paired t-test. Fear extinction was analyzed separately across the five sessions (Twelve extinction trials were averaged in bins of two trials per session). A polynomial linear trend test was used to assess within-session extinction for each extinction session, as well as across sessions, to evaluate the overall extinction trajectory. Paired t-tests were used to assess learning-related changes by comparing approach ratios between Session 1 and Session 6 in the appetitive conditioning paradigm. Additionally, paired t-tests were used to compare the number of nosepoke responses between the final rewarded session and subsequent punishment sessions in the positive reinforcement/positive punishment task.

### Fiber Photometry Recordings

ACh release in DLS was recorded by using a fiber photometry approach during different behavioral paradigms. Behavioral experiments were conducted in order from the least stressful to the most stressful, as follows: cue deliveries, appetitive conditioning, fear conditioning, and extinction. Reward learning and punishment were conducted in a different group of mice.

#### Fiber photometry general approach

Viral infusion and gross expression of ACh sensor iACh.Sn.FR (pAAV.hSynap.iACh.Sn.FR) in DLS neurons allowed us to quantify fluorescent signals and record through a permanently implanted fiberoptic (400μm fiber diameter, 0.48 NA) affixed to each mouse’s skull (described in surgical procedures). The fiber photometry recording system uses two light-emitting diodes (LED, Thorlabs; Lux RZ10X, TDT) controlled by an LED driver (Thorlabs; Lux RZ10X, TDT) at 465nM and 405nm (an isosbestic control channel). LED emissions pass through several filters and are reflected off a series of dichroic mirrors (Fluorescence MiniCube, Doric; LUX RZ10X, TDT), allowing emission and recording of the resulting excitation through the same optical system. LEDs were controlled by a real-time signal processor (RZ5P, RZ10X; Tucker-Davis Technologies), and emission signals from each LED stimulation were determined via multiplexing. The fluorescent signals were collected via a photoreceiver (Newport Visible Femtowatt

Photoreceiver Module, Doric). Synapse software (Tucker-Davis Technologies) was used to control the timing and intensity of the LEDs and to record the emitted fluorescent signals. The LED intensity was monitored via real-time fluorescence amplitude in Synapse software and maintained at a constant value between 350 mV and 500 mV across trials for each subject. For each event of interest (e.g., predictive cue, head entries, licks, shock), transistor-transistor logic (TTL) signals were used to timestamp onset times from Med-PC V software (Med Associates Inc.) and were detected via the RZ5P and RZ10X in the synapse software (explained in more detail in the analysis section below).

#### Fiber photometry analysis

The analysis of the fiber photometry data was conducted using a custom MATLAB pipeline, as described previously^4–7^. Raw 465nM (F465 channel) and isosbestic 405nM (F405 channel) traces were collected at a rate of 1000 samples per second (1kHz). The raw data from each individual channel (450nM or 465nM) were then minimally filtered using a lowess filter before calculating Δf/f values via polynomial curve fitting. For the lowess filter, the number of data points for calculating the filtered value was set to 0.0004 (values closer to 1 indicate stronger smoothing). Δf/f for the entire trace was calculated as (F465nm-F405nm)/F405nm. This transformation uses the isosbestic F405nm channel, which is not responsive to fluctuations in calcium, to control for calcium-independent fluctuations in the signal and to control for photobleaching. Then, the data from the resulting Δf/f trace were cropped around behavioral events using TTL pulses; for each experiment, 2s of pre-TTL and 18s of post-TTL Δf/f values were analyzed. Z-scores were calculated from the cropped trace by taking the pre-TTL Δf/f values as baseline (z-score = (TTLsignal - b_mean)/b_stdev, where the TTL signal is the Δf/f value for each post-TTL time point, b_mean is the baseline mean, and b_stdev is the baseline standard deviation). This allowed for the determination of calcium events that occurred at the precise moment of each significant behavioral event.

#### Statistical analyses of fiber photometry signals

For statistical analysis, peak height and area under the curve (AUC) values were calculated for each individual trace around identified behaviorally relevant events via trapezoidal numerical integration on each of the z-scores across a fixed timescale, which varied based on experiment. The AUC values were averaged within animals, and statistical comparisons were performed at the animal level. As with the behavioral analyses, we used paired t-tests to compare AUC values across learning (e.g., Session 1 vs. Session 6) within the appetitive conditioning paradigm. For the positive reinforcement/positive punishment experiment, we compared AUCs from the last rewarded session to each punishment session using paired t-tests. We also compared CS+-evoked AUC and shock-evoked AUC within animals using a paired t-test to assess how neural responses differentiated predictive cues from aversive outcomes. For the analysis of the fear extinction CS+ AUCs, we used a repeated-measures ANOVA followed by Dunnett’s post hoc tests to determine which sessions or conditions differed from EXT1 AUC. For the statistical analysis of the presentation of neural and valenced stimuli, we used independent sample t-tests (critical value = 0).

### Optogenetic Stimulation of DLS ACh release

Optogenetic stimulation was delivered via fiber-coupled diode laser system (589 nm, OptoEngine LLC, #FC-33-473-100) powered by a dedicated laser power supply (Model PSU-II-LED, OptoEngine). Timing and pulse parameters (i.e., frequency, pulse duration, train duration) were controlled by a Pulse Pal stimulator (Sanworks LLC). Laser intensity was daily measured before experiments using a power meter (PM100D, ThorLabs) to ensure that laser intensity was maintained constant across trials and experiments.

#### Validation of Optogenetic excitation using Fiber Photometry

Wild-type mice were infused in DLS with a viral compound consisting of rAAV-ChAT-Cre-WPRE-HGH polyAAV2/5 and pAAV-Syn-FLEX-rc[ChrimsonR-tdTomato] to infect only Cholinergic Interneurons (CINs) and acquire excitation in CINs via optogenetic photostimulation. Control animals received rAAV-ChAT-Cre-WPRE-HGH polyAAV2/5 and pAAV-hSyn-DIO-mCherry. The same animals were also injected with pAAV.hSynap.iACh.Sn.FR to allow us to record optogenetically stimulated instantaneous change in ACh release. A 400-μm fiber optic was implanted into the DLS. DLS ACh release change via optogenetic CIN excitation was validated with different laser intensities (589nm, 10sec, 10Hz, 2mW; 589nm, 10sec, 10Hz, 5mW; 589nm, 10sec, 10Hz, 8mW). Animals with no ACh.Sn.FR signal, and no ACh.Sn.FR increase during the optogenetic stimulations, were removed from the cohort.

#### Conditioned Place Aversion

Different laser intensities (589nm, 5sec, 10Hz, 2mW; 589nm, 5sec, 10Hz, 5mW; 589nm, 5sec, 10Hz, 8mW) were tested, and 5-second-long photostimulation was delivered with 5 5-second intervals in between stimulations in the test session when the test animal entered the preferred chamber.

#### Open Field Test

Animals were allowed to explore the open field for 5 minutes freely. Subsequent to the first trial, animals restart another 5-minute trial without an interval. On the second trial, animals received 30 seconds of photostimulation (589nm, 30sec, 10Hz, 8mW) with a 30-second interval in between stimulations.

#### Fear Conditioning/Extinction

Test was conducted according to the protocol stated under the behavior section, with a modification in the fear conditioning session. In order to achieve more robust fear extinction performance, we decreased the footshock intensity to 0.5 mA. Subsequently, the mice received only 3 extinction sessions instead of 5. Photostimulation (589nm, 10sec, 10Hz, 8mW) was delivered throughout fear conditioning and extinction sessions during each 10-second-long CS+.

#### Learning Curve Quantification

To determine whether optogenetically-evoked ACh release in the DLS influenced the acquisition and extinction of conditioned freezing behavior, freezing percentages were measured across trials during fear conditioning and extinction sessions. For each mouse, the trial-by-trial freezing data were fit using a standard exponential learning function:

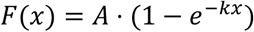

where F(x) is the freezing level on trial x, A is the asymptotic maximum freezing level, and k is the learning rate. This approach estimates how quickly freezing responses increased during conditioning or decreased during extinction. Curve fitting was performed for each mouse individually using nonlinear least-squares regression (lsqcurvefit in MATLAB). Parameter initialization was based on the maximum observed freezing value for A and a slope starting value of 0.5 for k. The learning rate parameter k was taken as the primary measure of learning speed. Learning rate values were then compared between the control group (mCherry) and the Chrimson group. Statistical comparison was performed using a Wilcoxon rank-sum test due to non-normal parameter distributions. In the fear conditioning session, Chrimson stimulation of cholinergic interneurons increased the learning rate, indicating faster acquisition of freezing. During extinction, the same stimulation reduced the rate of decline in freezing, indicating slower extinction. These effects show that optogenetically driven ACh release in the DLS facilitated the formation of fear associations and impeded their subsequent reduction.

#### Statistical analyses of the fiber photometry and behavioral measures during optogenetics

We used a repeated measures ANOVA (mCherry vs Chrimson [viral group] and 2mW vs 5mW vs 8mW laser [laser power]) followed by Sidak post-hocs to examine the AUCs during different optogenetic stimulation parameters. We used unpaired t-tests to compare the mCherry and Chrimson group learning rates during fear conditioning and fear extinction. We also used a repeated measures ANOVA to compare the mCherry and Chrimson group freezing levels during fear extinction.

### Simultaneous in vivo calcium imaging and optogenetic stimulation of DLS neural ensembles

#### Single-cell calcium imaging general approach

For calcium imaging at the single-cell level, we used endoscopic miniature microscopes (nVoke^TM^ miniature microscope, Inscopix), as described in previous publications. Calcium signals were recorded via the calcium sensor (AAV1.CaMK2a.GCaMP6m.WPRE.SV40). Signals were recorded on the miniature microscopes through the GRIN lens (0.5 mm diameter, 4.0 mm length, 0.5 NA; #1050-004417, Inscopix), allowing for the resolution of single cells. Single-cell activity in the DLS was recorded in awake and behaving animals. During each behavioral session, the miniscope was attached to the baseplate that was implanted previously (see surgical section for description). The imaging parameters (gain, LED power, and focus) were determined for each animal to ensure recording quality and were kept constant throughout the study. Data acquisition was conducted on Inscopix Data Acquisition Software (IDAS, v2.3.0, Inscopix), and sessions were video recorded using the nVision^TM^ System (Inscopix). At the end of the recording session, the miniscope was removed, and the baseplate cover was replaced.

#### Optogenetic manipulation within a single-cell imaging approach

Optogenetic stimulation was delivered via the nVoke^TM^ 2.0 system (Inscopix). Photostimulation intensity (OG-LED 620 ± 30nm, 10Hz, 5mW) was adjusted on Inscopix Data Acquisition Software (IDAS, v2.3.0, Inscopix) and maintained constant across trials and experiments. We used 5 mW LED stimulation for the combined imaging–optogenetics experiments because the nVoke system caps output at 5 mW, whereas the standalone optogenetics experiments employed 8–10 mW laser power. Timing and pulse parameters (i.e., frequency, pulse duration, train duration) were controlled by a Pulse Pal stimulator (Sanworks LLC).

#### Validation of Optogenetic excitation during in vivo calcium imaging

Wild-type mice were infused with DLS with a viral compound consisting of rAAV-ChAT-Cre-WPRE-HGH polyAAV2/5 and pAAV-Syn-FLEX-rc[ChrimsonR-tdTomato] to infect only Cholinergic Interneurons (CINs) and acquire excitation in CINs via optogenetic photostimulation. Control animals received rAAV-ChAT-Cre-WPRE-HGH polyAAV2/5 and pAAV-hSyn-DIO-mCherry. The same animals were also injected with AAV1.CaMK2a.GCaMP6m.WPRE.SV40 to allow us to record optogenetically stimulated instantaneous change in single-cell neuronal ensemble activity within the DLS population. The GRIN lens (0.5 mm diameter, 4.0 mm length, 0.5 NA; #1050-004417, Inscopix) was implanted in the DLS. DLS single-cell neuronal ensemble activity changes following optogenetic CIN excitation were validated using baseline recordings. Baseline recordings consisted of 10-second-long 4 photostimulation (OG-LED 620 ± 30nm, 10Hz, 5mW, 10sec) with randomized intervals (30sec, 45sec, 50sec) and calcium activity recordings of neuronal ensembles throughout the entire trial. Animals with no GCaMP signal were removed from the cohort.

#### Baseline recordings

In order to examine the effects of ACh stimulation at baseline levels, we recorded DLS single cell ensemble calcium dynamics while behaviorally animals received either 4 or 12 pseudorandom LED stimulations (590 nm, 10Hz, 5mW, 10sec) in MED PC operant chambers.

#### Fear Conditioning/Extinction

Test was conducted according to the protocol described for the optogenetics experiments above. Single-cell calcium recordings were carried out throughout the entire sessions, and photostimulation (LED 590 nm, 10Hz, 5mW, 10sec) was delivered throughout fear conditioning and extinction sessions during each 10-second-long CS+.

#### Image processing and signal extraction

Data was acquired at 20 frames per second using nVoke^TM^ miniature microscopes (Inscopix) via Inscopix Data Acquisition Software (IDAS, v2.3.0, Inscopix). Image processing was accomplished using Inscopix Data Processing Software (IDPS, v1.3.1, Inscopix). Raw videos were pre-processed by applying 2x spatial downsampling to reduce file size and processing time, and isolated dropped frames were corrected. No temporal downsampling was applied. Lateral movement was corrected by using a portion of a single reference frame using Inscopix Data Processing Software (IDPS, v1.3.1). Images were cropped to remove post-registration borders and sections in which cells were not observed. Videos were then exported as TIF stacks for analysis. After motion correction and cropping, a constrained non-negative matrix factorization algorithm optimized for micro-endoscopic imaging (CNMF-E^9^) was utilized to extract fluorescence traces from neurons. CNMF-E cell detection parameters were as follows: patch_dims = 50, 50; K = 20; gSiz = 20; gSig = 12; min_pnr = 20; min_corr = 0.8; max_tau = 0.400. Considering calcium fluctuations can exhibit negative transients, associated with a pause in firing 69, we did not constrain temporal components to >=0. The ΔF/F values were computed for the whole field of view as the output pixel value was represented as a relative percent change from the baseline. Raw CNMF-E traces were used for all analyses. The spatial mask and calcium time series of each cell were manually inspected using the IDPS interface. Cells found to be duplicated or misdetected due to neuropils or other artifacts were discarded.

#### Identification of cholinergic interneurons

In the Chrimson group, a subset of recorded neurons was expected to be DLS cholinergic interneurons (CINs), which express Chrimson and therefore exhibit reliable, time-locked calcium responses to optogenetic stimulation. To prevent these directly stimulated CINs from dominating the population activity structure, we identified and excluded them from all ensemble analyses.

For each neuron, z-scored calcium traces were aligned to the LED stimulation onset. CINs were classified based on three criteria reflecting time-locked and trial-consistent activation:

1. **Peak activation magnitude:** The maximum z-scored activity within the LED stimulation window (frames *n_pre+1: n_pre+200*) was required to exceed a threshold across all trials (z-score ≥ 1).
2. **Trial-to-trial response reliability:** We computed the average pairwise correlation of the LED-evoked response across trials, requiring a mean inter-trial correlation ≥ 0.3.
3. **Sustained activation profile:** To ensure activation was stimulus-evoked rather than spontaneous, the elevation in activity relative to trial baseline was required to fall below a sustained activation threshold (Δ < 15 z-units).

Cells meeting all three criteria were classified as CINs and removed from subsequent low-dimensional trajectory and population analysis. This procedure was performed using custom MATLAB code. To verify that results did not depend on the stringency of CIN exclusion, we also evaluated more liberal criteria (e.g., reduced inter-trial consistency threshold and increased sustained activation threshold), which resulted in a *larger* set of excluded neurons. Importantly, all population-level findings, including state-space trajectory geometry, distance-from-baseline dynamics, and group comparisons, were unchanged when using these more liberal thresholds. Thus, conclusions regarding non-CIN population dynamics were robust to the specific CIN classification criteria.

#### CIN Classification Validation

To ensure effective exclusion of CINs, we trained a random-forest classifier on feature vectors derived from identified CINs and non-CINs, including mean, variance, range, skewness, kurtosis, event rate, event amplitude, lock index, and latency jitter. The classifier was then applied to both baseline and fear-conditioning recordings. Out-of-bag accuracy exceeded 98%, demonstrating near-perfect separability between CINs and non-CINs. A conservative CIN-likeness probability threshold of 0.2—corresponding to the empirical minimum between CIN and non-CIN probability distributions—was used to detect any residual CIN-like activity. In the baseline recordings, only 2 of 513 Chrimson cells (0.39%) and 1 of 311 mCherry cells (0.32%) exceeded this threshold. During fear conditioning, only 1 of 958 Chrimson cells (0.10%) and none of 462 mCherry cells (0.00%) were classified as CIN-like. These results confirm that both datasets were overwhelmingly composed of non-cholinergic neurons.

#### Calcium activity quantification around behavioral events

Stimulus-evoked activity was assessed by aligning calcium activity traces around the onset of the cue or stimulus. Transistor-transistor logic (TTL) signals from MedPC were directly fed to the nVoke system. The behavioral apparatus was interfaced with the miniature microscope via BNC cables, and the onset of behavioral events - cues and shocks - was associated with a frame of the video using TTLs. The Δf/f values for each movie frame were calculated as M’(x,y,t) = (M(x,y,t)-Fbaseline(x,y))/Fbaseline(x,y) where M’ is the output movie with Δf/f values, M(x,y,t) is the value for the pixel coordinate (x,y) at the t frame of the movie, and Fbaseline(x,y) is the baseline value for the (x,y) coordinate. The raw Δf/f data from the CNMF-E program were exported and used for peri-event analysis^9^. Data was cropped around each significant event (cue presentations; TTL) and z-scored to normalize for baseline differences. Z-scores were calculated by taking the pre-TTL Δf/f values as baseline (z-score = (TTLsignal - b_mean)/b_stdev, where TTL signal is the Δf/f value for each post-TTL time point, b_mean is the baseline mean, and b_stdev is the baseline standard deviation). For traces aligned around a stimulus, a baseline window of 2 seconds prior to stimulus onset was used. Z-scored traces were then averaged across trials to create one trace per neuron for each stimulus type. To quantify the magnitude of the response, peak height (the maximum value of the calcium transient) and area under the curve (AUC) for each animal’s averaged population trace were calculated. Values across a 10-second window beginning at stimulus onset were averaged, and this value was used to determine the response profile of each individual cell.

#### Classification of Non-CIN Neuronal Responses to the CS+

Following the identification and removal of cholinergic interneurons (CINs) that were directly responsive to Chrimson stimulation, the remaining neurons were classified as non-CIN cells. These neurons did not receive direct optical activation; therefore, any modulation in their activity during behavior reflects network-level effects of optogenetically-evoked acetylcholine release, rather than direct stimulation. For each non-CIN neuron, z-scored calcium traces were aligned to the onset of the CS+ and averaged across trials. We quantified mean activity during the CS+ presentation window and during the subsequent post-CS+ window. Neurons whose average activity during the CS+ increased above 2.5 z-score units were classified as *Excited*, whereas neurons whose activity fell below −0.8 z-score units during the CS+ were classified as *Inhibited*. A distinct group of neurons did not show modulation during the CS+ itself but exhibited a delayed increase in activity immediately after stimulus offset. These neurons were classified as *Rebound*-responsive and were defined by low activity during the CS+, combined with a positive shift in activity in the post-stimulus window. Neurons that did not meet any of these criteria were considered *Unchanged*.

#### Hierarchical clustering analysis

Using calcium traces from all non-CIN cells detected in baseline sessions, we employed a hierarchical clustering approach to group cells based on their responses to ACh stimulation. We used the “clustergram” Matlab function and the “correlation” distance metric to group the cell activity. We then combined all the cells that were clustered in 3 main nodes to visualize the group characteristics.

#### State-space analysis

To examine the evolution of population activity dynamics during learning, we quantified neural state-space trajectories from single-cell calcium imaging data. For each mouse, ΔF/F traces were extracted from individual non-CIN neurons across repeated trials of the behavioral session. For analysis, we focused on a defined epoch encompassing both the conditioned stimulus (CS+) and unconditioned stimulus (US) periods (frames 41–340; equivalent to 15 seconds after the CS+ onset). For each neuron, data were organized into a three-dimensional array (*time × neurons × trials*). Trial-averaged population activity was then computed to obtain a mean ensemble activity pattern across time. Principal component analysis (PCA) was applied to the trial-averaged time-by-neuron matrix using MATLAB’s `pca` function, and the first three principal components (PCs) were retained to visualize the dominant low-dimensional population dynamics. These three PCs captured the majority of explainable variance in population activity (typically ∼45–85% across groups). The resulting reduced trajectory (PC1–PC3) was plotted as a continuous three-dimensional neural state-space trajectory, color-coded by elapsed time. To quantify the magnitude of population state shifts during the behavioral epoch, we computed the Euclidean distance between the ensemble state at each time point and a baseline reference state, defined as the mean state during the first 10 frames of the analysis window. Additional metrics included (1) **trajectory length** (the cumulative Euclidean path length through state space), (2) **maximum distance from baseline,** (3) **mean distance across time,** and (4) **area under the distance-time curve (AUC).** These measures provide complementary estimates of how far and how dynamically the neural population diverged from its baseline representational state during the session. The analysis was performed identically for mice expressing **Chrimson** in dorsolateral striatal cholinergic interneurons (DLS CINs) and for **mCherry** control mice during both the fear conditioning session (FC) and the last session of fear extinction (EXT3). Group comparisons were conducted on trajectory metrics to test whether optogenetic stimulation altered the geometry or magnitude of striatal population state transitions during learning.

#### Analysis of baseline calcium dynamics to predict ACh-evoked response polarity

Single-cell calcium traces were obtained from Inscopix nVoke recordings sampled at 20 Hz. Each column corresponded to an identified non-CIN neuron, and each row corresponded to a time point. Columns consisting entirely of NaNs or without valid identifiers were removed. For each neuron, we extracted the full continuous activity trace (“whole trace”) as well as an LED-free trace. LED onset times were determined from mouse-specific TTL files, and all frames within −1/+10 seconds of each LED event were removed to generate a spontaneous-only baseline trace. We computed a comprehensive set of descriptive statistics for each neuron using the full recording: mean activity, standard deviation, maximum, minimum, range, skewness, kurtosis, spontaneous event rate, and mean event amplitude. Spontaneous calcium events were defined as local maxima exceeding one standard deviation above the trace mean. All features were compiled along with the mouse ID, cell ID, and CIN-evoked response classification (“Excited” or “Inhibited”). To test whether spontaneous activity predicted response polarity, we extracted an analogous set of features exclusively from LED-free traces. These baseline features included mean activity, variance, minimum, maximum, range, skewness, kurtosis, event rate, and event amplitude. Features were computed after demeaning or z-scoring when appropriate. Missing values were imputed using column-wise means.

#### GLM Classification of Excited vs Inhibited Cells

A binomial generalized linear model (GLM; logistic regression) was used to predict whether each neuron became excited or inhibited during CIN stimulation based solely on baseline features. The model followed the equation:

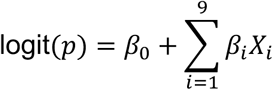

where Xᵢ were the nine baseline features. Model performance was quantified using overall prediction accuracy, the chi-square statistic for improvement over the null model, and statistical significance of individual coefficients

#### Random Forest Classification and Permutation Importance

To assess nonlinear and multivariate interactions among features, we trained a Random Forest classifier (MATLAB TreeBagger; 500 trees, out-of-bag prediction enabled). Out-of-bag accuracy provided an unbiased estimate of classification performance. Feature importance was assessed using permutation importance, in which individual features were randomly shuffled to measure their impact on out-of-bag prediction error.

#### Statistical analyses for the behavioral and single-cell calcium data

We quantified learning-related activity changes during fear conditioning and extinction in the single-cell imaging/optogenetics cohort as described above. To assess the impact of optogenetic stimulation, we compared single-cell AUC values between pre-LED and LED epochs, as well as between mCherry and Chrimson groups. For learning-related changes, AUCs for each cell during fear conditioning (averaged across the 6 CS+ trials) and extinction session 3 (averaged across the 12 CS+ trials) were compared using paired t-tests (equal SD assumed) for within-subject comparisons and using Welch’s t-test (equal SD not assumed) for between-group comparisons. In addition, mean distances from baseline activity (averaged per animal) were compared across experimental conditions using Welch’s t-tests to account for unequal variances.

## Notes

### Competing Interest Statement

The authors have declared no competing interest.

## References

1. Fujii, S., Ji, Z. & Sumikawa, K. Inactivation of α7 ACh receptors and activation of non-α7 ACh receptors both contribute to long term potentiation induction in the hippocampal CA1 region. Neuroscience Letters 286, 134–138 (2000).

2. Nakauchi, S. & Sumikawa, K. Endogenously released ACh and exogenous nicotine differentially facilitate long-term potentiation induction in the hippocampal CA1 region of mice. European Journal of Neuroscience 35, 1381–1395 (2012).

3. Newman, E. L., Gupta, K., Climer, J. R., Monaghan, C. K. & Hasselmo, M. E. Cholinergic modulation of cognitive processing: insights drawn from computational models. Front. Behav. Neurosci. 6, (2012).

4. Hasselmo, M. E. The role of acetylcholine in learning and memory. Current Opinion in Neurobiology 16, 710–715 (2006).

5. Sarter, M., Bruno, J. P. & Givens, B. Attentional functions of cortical cholinergic inputs: What does it mean for learning and memory? Neurobiology of Learning and Memory 80, 245–256 (2003).

6. Klinkenberg, I., Sambeth, A. & Blokland, A. Acetylcholine and attention. Behavioural Brain Research 221, 430–442 (2011).

7. Howe, W. M. et al. Prefrontal Cholinergic Mechanisms Instigating Shifts from Monitoring for Cues to Cue-Guided Performance: Converging Electrochemical and fMRI Evidence from Rats and Humans. J. Neurosci. 33, 8742–8752 (2013).

8. Collins, A. L. et al. Nucleus Accumbens Cholinergic Interneurons Oppose Cue-Motivated Behavior. Biol Psychiatry 86, 388–396 (2019).

9. Mohebi, A., Collins, V. L. & Berke, J. D. Accumbens cholinergic interneurons dynamically promote dopamine release and enable motivation. eLife 12, e85011 (2023).

10. Pérez-González, D., Lao-Rodríguez, A. B., Aedo-Sánchez, C. & Malmierca, M. S. Acetylcholine modulates the precision of prediction error in the auditory cortex. eLife 12, RP91475 (2024).

11. Sturgill, J. F. et al. Basal forebrain-derived acetylcholine encodes valence-free reinforcement prediction error. 2020.02.17.953141 Preprint at 10.1101/2020.02.17.953141 (2020).

12. Robert, B. et al. A functional topography within the cholinergic basal forebrain for encoding sensory cues and behavioral reinforcement outcomes. eLife 10, e69514 (2021).

13. Gritton, H. J. et al. Cortical cholinergic signaling controls the detection of cues. Proceedings of the National Academy of Sciences 113, E1089–E1097 (2016).

14. Zhang, Z. et al. Accumbal acetylcholine signals associative salience during learning. bioRxiv 2025.01.06.631529 (2025) doi:10.1101/2025.01.06.631529.

15. Disney, A. A. & Higley, M. J. Diverse Spatiotemporal Scales of Cholinergic Signaling in the Neocortex. J Neurosci 40, 720–725 (2020).

16. Dautan, D. et al. A Major External Source of Cholinergic Innervation of the Striatum and Nucleus Accumbens Originates in the Brainstem. J. Neurosci. 34, 4509–4518 (2014).

17. Lim, S. A. O., Kang, U. J. & McGehee, D. S. Striatal cholinergic interneuron regulation and circuit effects. Front Synaptic Neurosci 6, 22 (2014).

18. Woolf, N. J. & Butcher, L. L. Cholinergic neurons in the caudate-putamen complex proper are intrinsically organized: A combined evans blue and acetylcholinesterase analysis. Brain Research Bulletin 7, 487–507 (1981).

19. Threlfell, S. et al. Striatal Dopamine Release Is Triggered by Synchronized Activity in Cholinergic Interneurons. Neuron 75, 58–64 (2012).

20. Nelson, A. B. et al. Striatal Cholinergic Interneurons Drive GABA Release from Dopamine Terminals. Neuron 82, 63–70 (2014).

21. Zucca, S., Zucca, A., Nakano, T., Aoki, S. & Wickens, J. Pauses in cholinergic interneuron firing exert an inhibitory control on striatal output in vivo. Elife 7, e32510 (2018).

22. Maurice, N. et al. Striatal Cholinergic Interneurons Control Motor Behavior and Basal Ganglia Function in Experimental Parkinsonism. Cell Rep 13, 657–666 (2015).

23. Howe, M. et al. Coordination of rapid cholinergic and dopaminergic signaling in striatum during spontaneous movement. eLife 8, e44903 (2019).

24. Apicella, P., Ravel, S., Deffains, M. & Legallet, E. The role of striatal tonically active neurons in reward prediction error signaling during instrumental task performance. J Neurosci 31, 1507–1515 (2011).

25. Bouabid, S. et al. Distinct spatially organized striatum-wide acetylcholine dynamics for the learning and extinction of Pavlovian associations. Nat Commun 16, 5169 (2025).

26. Huang, Z. et al. Dynamic responses of striatal cholinergic interneurons control behavioral flexibility. Sci Adv 10, eadn2446.

27. Balleine, B. W., Delgado, M. R. & Hikosaka, O. The Role of the Dorsal Striatum in Reward and Decision-Making. J. Neurosci. 27, 8161–8165 (2007).

28. Featherstone, R. E. & McDonald, R. J. Dorsal striatum and stimulus-response learning: lesions of the dorsolateral, but not dorsomedial, striatum impair acquisition of a stimulus-response-based instrumental discrimination task, while sparing conditioned place preference learning. Neuroscience 124, 23–31 (2004).

29. Yin, H. H., Ostlund, S. B., Knowlton, B. J. & Balleine, B. W. The role of the dorsomedial striatum in instrumental conditioning. European Journal of Neuroscience 22, 513–523 (2005).

30. Devan, B. D., Hong, N. S. & McDonald, R. J. Parallel associative processing in the dorsal striatum: Segregation of stimulus–response and cognitive control subregions. Neurobiology of Learning and Memory 96, 95–120 (2011).

31. Stanley, A. T., Lippiello, P., Sulzer, D. & Miniaci, M. C. Roles for the Dorsal Striatum in Aversive Behavior. Front Cell Neurosci 15, 634493 (2021).

32. Delgado, M. R., Li, J., Schiller, D. & Phelps, E. A. The role of the striatum in aversive learning and aversive prediction errors. Philos Trans R Soc Lond B Biol Sci 363, 3787–3800 (2008).

33. Alloway, K. D., Smith, J. B., Mowery, T. M. & Watson, G. D. R. Sensory Processing in the Dorsolateral Striatum: The Contribution of Thalamostriatal Pathways. Front. Syst. Neurosci. 11, (2017).

34. Graybiel, A. M. & Grafton, S. T. The Striatum: Where Skills and Habits Meet. Cold Spring Harb Perspect Biol 7, a021691 (2015).

35. Malvaez, M. & Wassum, K. M. Regulation of habit formation in the dorsal striatum. Current Opinion in Behavioral Sciences 20, 67–74 (2018).

36. Borden, P. M. et al. A fast genetically encoded fluorescent sensor for faithful in vivo acetylcholine detection in mice, fish, worms and flies. 2020.02.07.939504 Preprint at 10.1101/2020.02.07.939504 (2020).

37. Picciotto, M. R., Higley, M. J. & Mineur, Y. S. Acetylcholine as a neuromodulator: cholinergic signaling shapes nervous system function and behavior. Neuron 76, 116–129 (2012).

38. Haam, J. & Yakel, J. L. Cholinergic modulation of the hippocampal region and memory function. Journal of Neurochemistry 142, 111–121 (2017).

39. Luchicchi, A., Bloem, B., Viaña, J. N. M., Mansvelder, H. D. & Role, L. W. Illuminating the role of cholinergic signaling in circuits of attention and emotionally salient behaviors. Front. Synaptic Neurosci. 6, (2014).

40. Teles-Grilo Ruivo, L. M., et al. Coordinated Acetylcholine Release in Prefrontal Cortex and Hippocampus Is Associated with Arousal and Reward on Distinct Timescales. Cell Rep 18, 905–917 (2017).

41. Abdulla, Z. I. et al. Acetylcholine signaling in the medial prefrontal cortex mediates the ability to learn an active avoidance response following learned helplessness training. bioRxiv 2023.09.23.559126 (2023) doi:10.1101/2023.09.23.559126.

42. Mineur, Y. S. et al. Hippocampal acetylcholine modulates stress-related behaviors independent of specific cholinergic inputs. Mol Psychiatry 27, 1829–1838 (2022).

43. Mineur, Y. S. et al. ACh signaling modulates activity of the GABAergic signaling network in the basolateral amygdala and behavior in stress-relevant paradigms. Mol Psychiatry 27, 4918–4927 (2022).

44. Tu, G., Wen, P., Halawa, A. & Takehara-Nishiuchi, K. Acetylcholine modulates prefrontal outcome coding during threat learning under uncertainty. eLife 13, RP102986 (2025).

45. Rajebhosale, P. et al. Functionally refined encoding of threat memory by distinct populations of basal forebrain cholinergic projection neurons. eLife 13, e86581 (2024).

46. Tu, G., Halawa, A., Yu, X., Gillman, S. & Takehara-Nishiuchi, K. Outcome-Locked Cholinergic Signaling Suppresses Prefrontal Encoding of Stimulus Associations. J. Neurosci. 42, 4202–4214 (2022).

47. Giovanniello, J. R. et al. A dual-pathway architecture for stress to disrupt agency and promote habit. Nature 640, 722–731 (2025).

48. Calabresi, P., Centonze, D., Gubellini, P., Pisani, A. & Bernardi, G. Acetylcholine-mediated modulation of striatal function. Trends in Neurosciences 23, 120–126 (2000).

49. Malenka, R. C. & Kocsis, J. D. Presynaptic actions of carbachol and adenosine on corticostriatal synaptic transmission studied in vitro. J. Neurosci. 8, 3750–3756 (1988).

50. Pakhotin, P. & Bracci, E. Cholinergic Interneurons Control the Excitatory Input to the Striatum. J. Neurosci. 27, 391–400 (2007).

51. Oldenburg, I. A. & Ding, J. B. Cholinergic modulation of synaptic integration and dendritic excitability in the striatum. Curr Opin Neurobiol 21, 425–432 (2011).

52. Pancani, T. et al. M4 mAChR-Mediated Modulation of Glutamatergic Transmission at Corticostriatal Synapses. ACS Chem Neurosci 5, 318–324 (2014).

53. Bernard, V., Normand, E. & Bloch, B. Phenotypical characterization of the rat striatal neurons expressing muscarinic receptor genes. J. Neurosci. 12, 3591–3600 (1992).

54. Sugita, S., Uchimura, N., Jiang, Z. G. & North, R. A. Distinct muscarinic receptors inhibit release of gamma-aminobutyric acid and excitatory amino acids in mammalian brain. Proceedings of the National Academy of Sciences 88, 2608–2611 (1991).

55. English, D. F. et al. GABAergic circuits mediate the reinforcement-related signals of striatal cholinergic interneurons. Nat Neurosci 15, 123–130 (2011).

56. Koós, T. & Tepper, J. M. Dual cholinergic control of fast-spiking interneurons in the neostriatum. J Neurosci 22, 529–535 (2002).

57. Rice, M. E. & Cragg, S. J. Nicotine amplifies reward-related dopamine signals in striatum. Nat Neurosci 7, 583–584 (2004).

58. Matityahu, L. et al. Acetylcholine waves and dopamine release in the striatum. Nat Commun 14, 6852 (2023).

59. Exley, R., McIntosh, J. M., Marks, M. J., Maskos, U. & Cragg, S. J. Striatal α5 nicotinic receptor subunit regulates dopamine transmission in dorsal striatum. J Neurosci 32, 2352–2356 (2012).

60. Campos, F., Alfonso, M. & Durán, R. In vivo modulation of α7 nicotinic receptors on striatal glutamate release induced by anatoxin-A. Neurochemistry International 56, 850–855 (2010).

61. Carpenedo, R. et al. Presynaptic kynurenate-sensitive receptors inhibit glutamate release. European Journal of Neuroscience 13, 2141–2147 (2001).

62. Beggiato, S. et al. Kynurenic acid, by targeting α7 nicotinic acetylcholine receptors, modulates extracellular GABA levels in the rat striatum in vivo. European Journal of Neuroscience 37, 1470–1477 (2013).

63. Gould, T. J. & Wehner, J. M. Nicotine enhancement of contextual fear conditioning. Behavioural Brain Research 102, 31–39 (1999).

64. Gould, T. J., Feiro, O. & Moore, D. Nicotine enhances trace cued fear conditioning but not delay cued fear conditioning in C57BL/6 mice. Behavioural Brain Research 155, 167–173 (2004).

65. Tian, S. et al. Nicotine enhances contextual fear memory reconsolidation in rats. Neurosci Lett 487, 368–371 (2011).

66. Kutlu, M. G. & Gould, T. J. Acute nicotine delays extinction of contextual fear in mice. Behavioural Brain Research 263, 133–137 (2014).

67. Kutlu, M. G. et al. Nicotine modulates contextual fear extinction through changes in ventral hippocampal GABAergic function. Neuropharmacology 141, 192–200 (2018).

68. Kutlu, M. G., Oliver, C., Huang, P., Liu-Chen, L.-Y. & Gould, T. J. Impairment of contextual fear extinction by chronic nicotine and withdrawal from chronic nicotine is associated with hippocampal nAChR upregulation. Neuropharmacology 109, 341–348 (2016).

69. Kutlu, M. G., Holliday, E. & Gould, T. J. High-affinity α4β2 nicotinic receptors mediate the impairing effects of acute nicotine on contextual fear extinction. Neurobiology of Learning and Memory 128, 17–22 (2016).

70. Kutlu, M. G., Tumolo, J. M., Holliday, E., Garrett, B. & Gould, T. J. Acute nicotine enhances spontaneous recovery of contextual fear and changes c-fos early gene expression in infralimbic cortex, hippocampus, and amygdala. Learn. Mem. 23, 405–414 (2016).

71. Lv, X. et al. M1 muscarinic activation induces long-lasting increase in intrinsic excitability of striatal projection neurons. Neuropharmacology 118, 209–222 (2017).

72. Bernard, V., Laribi, O., Levey, A. I. & Bloch, B. Subcellular Redistribution of m2 Muscarinic Acetylcholine Receptors in Striatal Interneurons In Vivo after Acute Cholinergic Stimulation. J. Neurosci. 18, 10207–10218 (1998).

73. Hersch, S. M., Gutekunst, C. A., Rees, H. D., Heilman, C. J. & Levey, A. I. Distribution of m1-m4 muscarinic receptor proteins in the rat striatum: light and electron microscopic immunocytochemistry using subtype-specific antibodies. J. Neurosci. 14, 3351–3363 (1994).

74. Abudukeyoumu, N., Hernandez-Flores, T., Garcia-Munoz, M. & Arbuthnott, G. W. Cholinergic modulation of striatal microcircuits. European Journal of Neuroscience 49, 604–622 (2019).

75. Jeon, J. et al. A Subpopulation of Neuronal M4 Muscarinic Acetylcholine Receptors Plays a Critical Role in Modulating Dopamine-Dependent Behaviors. J. Neurosci. 30, 2396–2405 (2010).

76. Shen, W. et al. M4 muscarinic receptor signaling ameliorates striatal plasticity deficits in models of L-DOPA-induced dyskinesia. Neuron 88, 762–773 (2015).

## References

1. Mesulam, M.-M., Mufson, E. J., Levey, A. I. & Wainer, B. H. Cholinergic innervation of cortex by the basal forebrain: Cytochemistry and cortical connections of the septal area, diagonal band nuclei, nucleus basalis (Substantia innominata), and hypothalamus in the rhesus monkey. J. Comp. Neurol. 214, 170–197 (1983).

2. Mesulam, M.-M., Mufson, E. J., Wainer, B. H. & Levey, A. I. Central cholinergic pathways in the rat: An overview based on an alternative nomenclature (Ch1–Ch6). Neuroscience 10, 1185–1201 (1983).

3. Kutlu, M. G., Ortega, L. A. & Gould, T. J. Strain-dependent performance in nicotine-induced conditioned place preference. Behav. Neurosci. 129, 37–41 (2015).

4. Kutlu, M. G. et al. Dopamine release in the nucleus accumbens core signals perceived saliency. Curr. Biol. 31, 4748–4761.e8 (2021).

5. Kutlu, M. G. et al. Dopamine signaling in the nucleus accumbens core mediates latent inhibition. Nat. Neurosci. 25, 1071–1081 (2022).

6. Kutlu, M. G., Tat, J., Christensen, B. A., Zachry, J. E. & Calipari, E. S. Dopamine release at the time of a predicted aversive outcome causally controls the trajectory and expression of conditioned behavior. Cell Rep. 42, 112948 (2023).

7. Zachry, J. E. et al. D1 and D2 medium spiny neurons in the nucleus accumbens core have distinct and valence-independent roles in learning. Neuron 112, 835–849.e7 (2024).

8. Dinckol, O., Wenger, N. H., Zachry, J. E. & Kutlu, M. G. Nucleus accumbens core single cell ensembles bidirectionally respond to experienced versus observed aversive events. Sci. Rep. 13, 22602 (2023).

9. Zhou, P. et al. Efficient and accurate extraction of in vivo calcium signals from microendoscopic video data. eLife 7, e28728 (2018).

