## Supplementary material for "Dorsolateral striatal acetylcholine reorganizes neural ensembles to anticipate threat": Supp figures 1-12

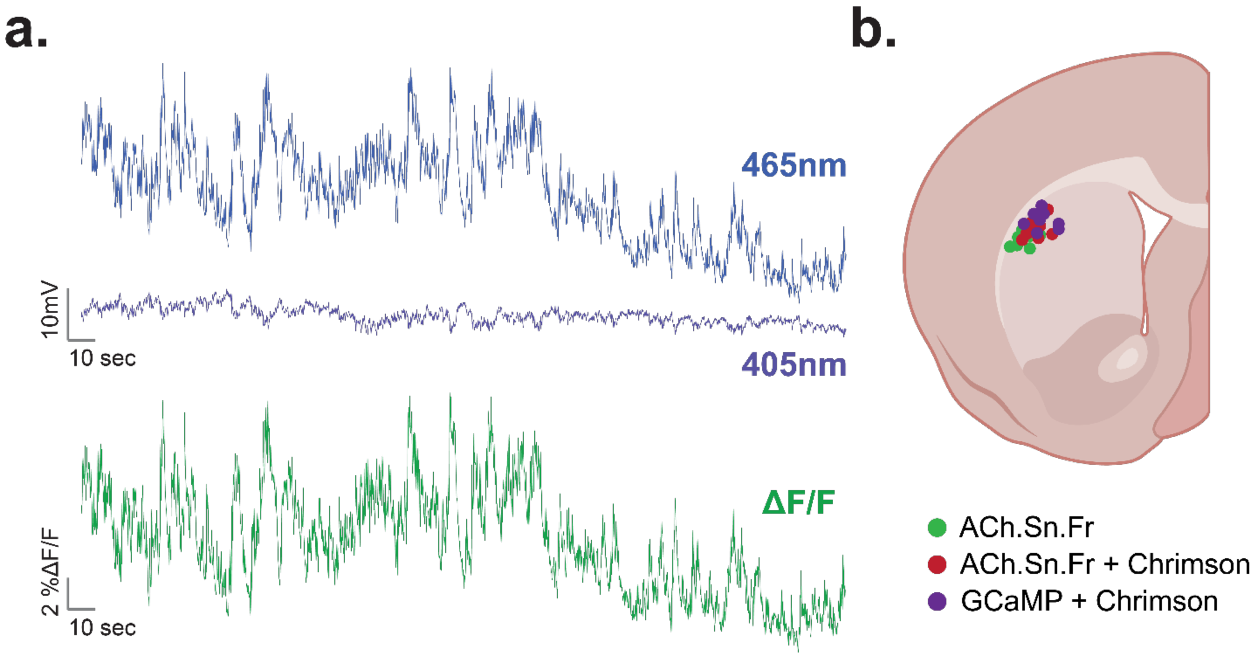


**Extended Data Figure 1. Validation of photometry signal acquisition and fiber placement in DLS.**

**(a)** Representative recording shows ACh release via GFP excitation using a 465nm LED. The change in fluorescence was divided by the baseline fluorescence, visualized via 405nm LED excitation, to calculate the signal-to-noise ratio as ΔF/F.

**(b)** Graphic representing validated implant placement in ACh.Sn.Fr mice, ACh.Sn.Fr + Chrimson mice (red), and GCaMP + Chrimson mice (purple).

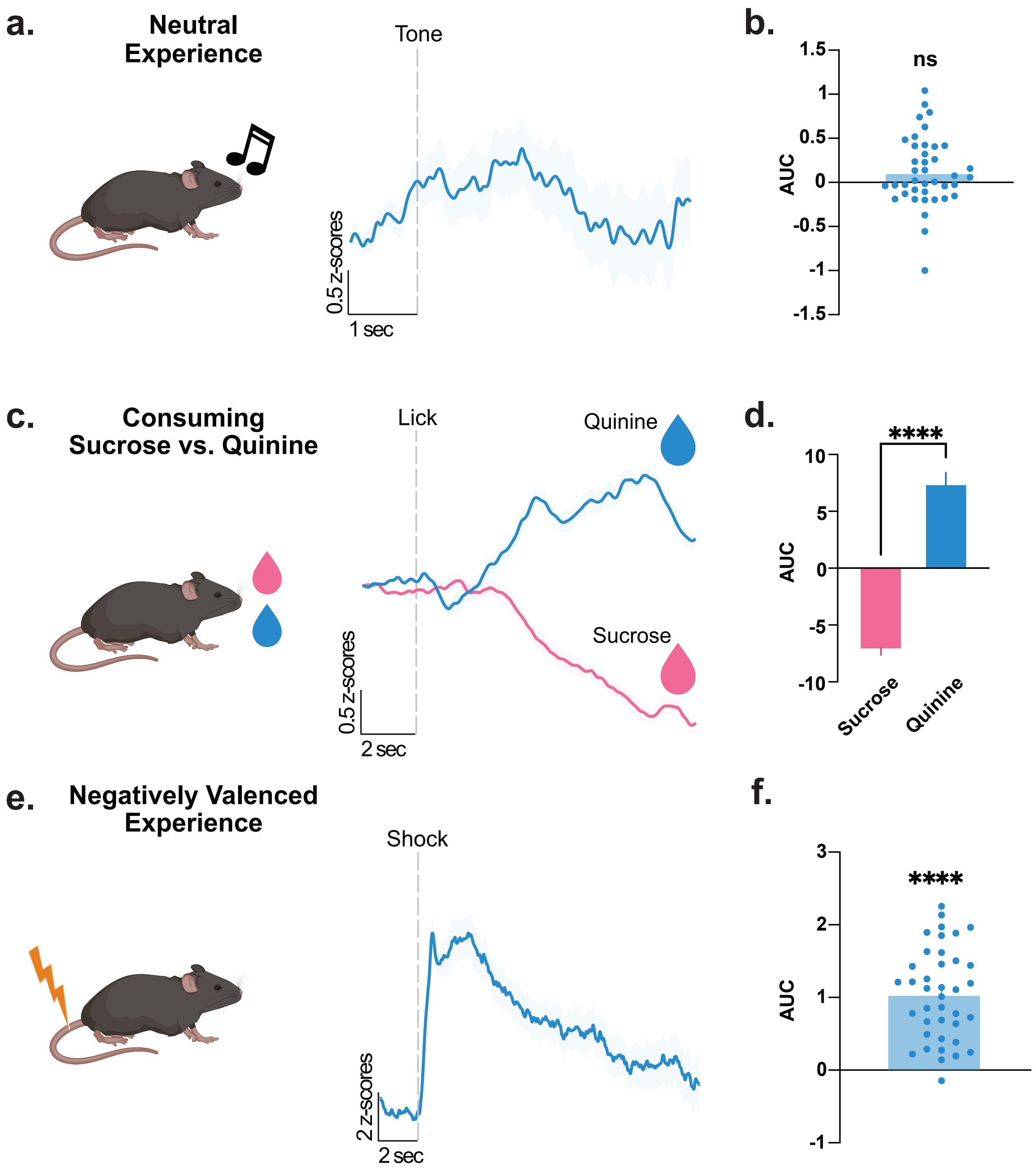


**Extended Data Figure 2. DLS acetylcholine responses to neutral, appetitive, and aversive stimuli.**

**(a-b)** DLS ACh release at the time of tone presentation as a neutral stimulus. No change in ACh release was observed at the time of tone presentation (One-sample t-test; $t(39)=1.840$, $p=0.0734$; n=40 tone presentations).

**(c-d)** DLS ACh release as mice consume either a 30% sucrose solution or a 0.32 g/L quinine solution via a retractable sipper. DLS ACh is significantly decreased at the time of a sucrose lick compared to a quinine lick (Unpaired t-test; $t(1307)=11.71$, $p<0.0001$; n=339-1000 lick events).

**(e-f)** DLS ACh release at the time of 1.0 mA footshock presentation as a negatively valenced stimulus. DLS ACh release is significantly increased at the time of footshock presentation (One-sample t-test; $t(39)=10.63$, $p<0.0001$; n=40 footshock presentations). Data represented as mean ± S.E.M. **** p < 0.0001, ns= not significant.

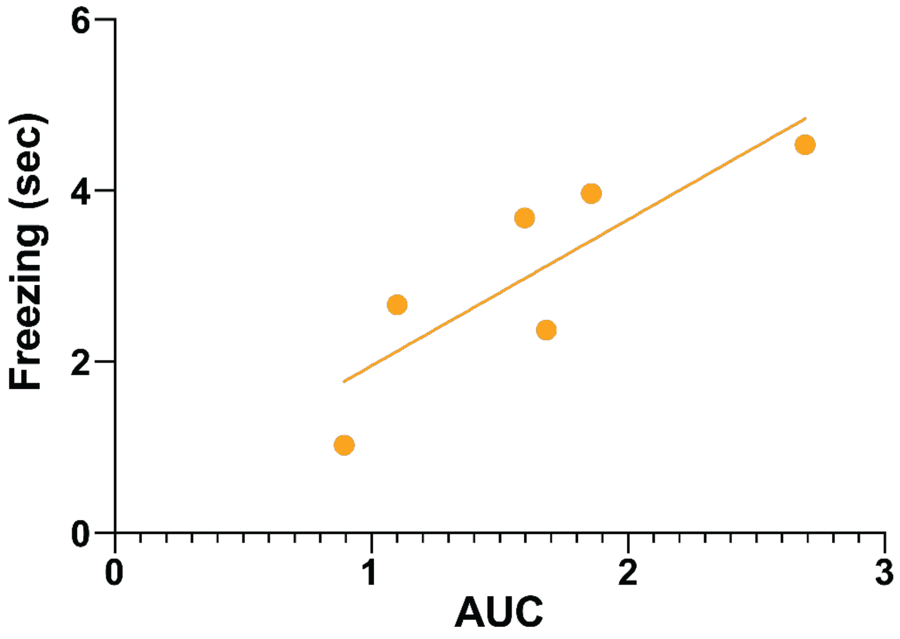


**Extended Data Figure 3. Positive relationship between DLS ACh signal magnitude and freezing behavior during fear conditioning.**
AUC values of DLS acetylcholine release during CS+ presentation on the fear conditioning session were positively correlated with freezing duration across trials (r=0.8471; *p*= 0.0333; n=8 mice). Each point represents a single CS+ trial, and the regression line indicates that larger ACh responses were associated with greater freezing behavior, reflecting stronger threat expression.

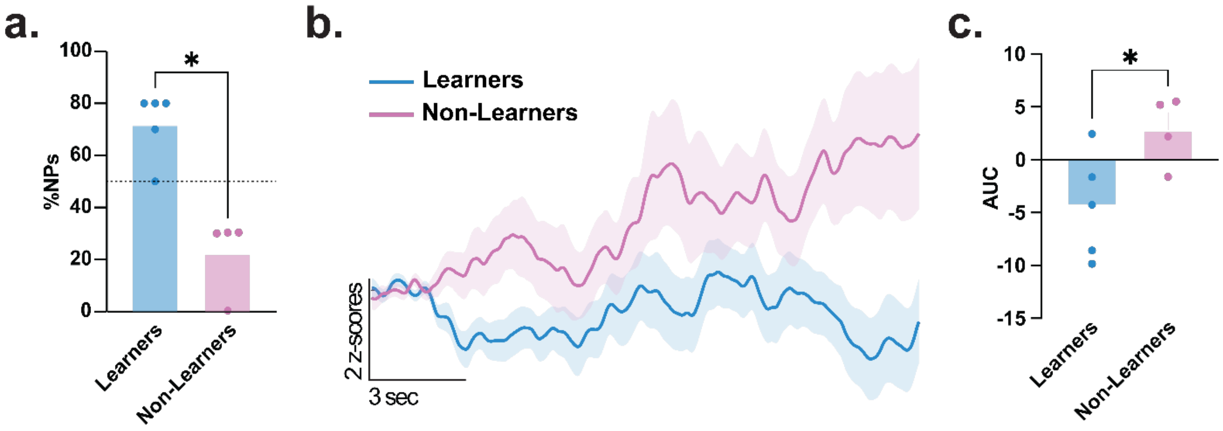


**Extended Data Figure 4. Behavioral performance and baseline DLS ACh differences between learner and non-learner mice during positive reinforcement.**

**(a)** Comparison of nosepoke percentage in learners and non-learners mice (n=4 mice, 2 males and 2 females) during the positive reinforcement session, with learner mice (n=5 mice, 3 males and 2 females) correctly nosepoking during the cue more frequently than non-learners (Unpaired t-test; t(7)=5.304, *p*=0.0011; n=4-5 mice per condition).

**(b)** Baseline DLS ACh release in learner and non-learner mice.

**(c)** Learner mice had significantly lower baseline DLS ACh release compared to non-learners (Unpaired t-test; t(7)=2.448, *p*=0.0442; n=4-5 mice per condition). Data represented as mean ± S.E.M. * $p < 0.05$.

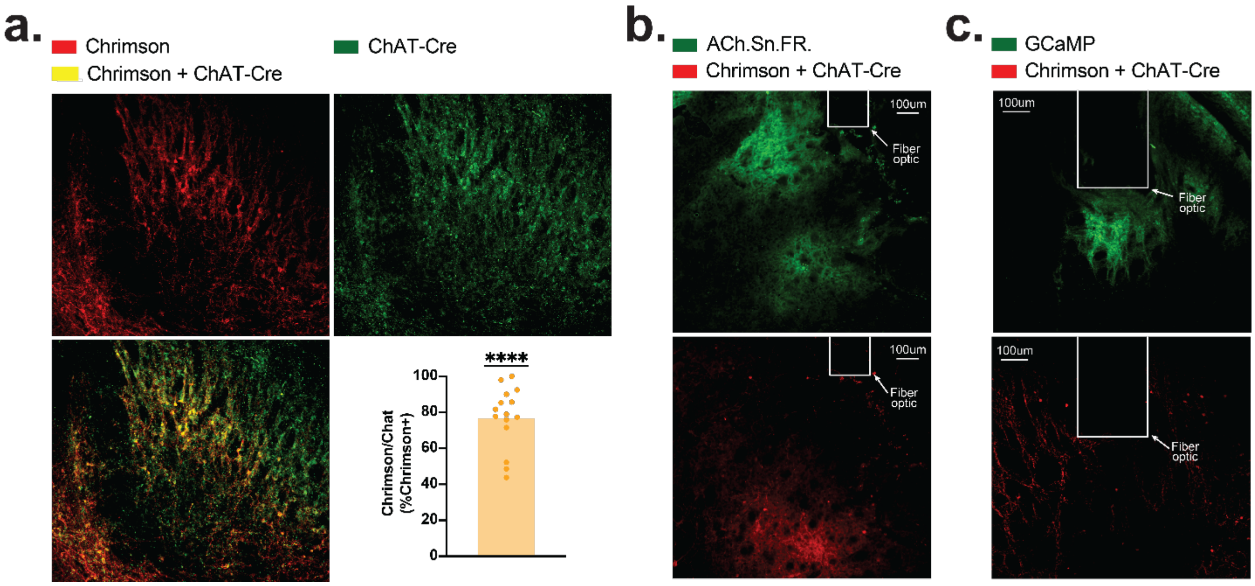


**Extended Data Figure 5. Verification of Cre-dependent Chrimson targeting to cholinergic interneurons.**

**(a)** Representative histology of basal forebrain neurons (M/L -1.5mm, A/P -0.34mm, D/V -5.3mm) following intracranial injection of AAV5-Syn-FLEX-rc[ChrimsonR-tdTomato] (red), AAVs AAV2/5-ChAT-Cre-WPRE-HGH (green), with co-infected neurons appearing yellow (One-sample t-test; $t(14)=6.170$, $p<0.0001$).

**(b)** Representative histology of DLS neurons following fiber optic implantation and intracranial injection of AAVs: AAV1.hSynap.iAChSnFR (green), AAV2/5-ChAT-Cre-WPRE-HGH, and AAV5-Syn-FLEX-rc[ChrimsonR-tdTomato] (red).

**(c)** Representative histology of DLS neurons following GRIN lens implantation and intracranial injection of AAVs: AAV1.CaMK2a.GCaMP6m.WPRE.SV40 (green), AAV2/5-ChAT-Cre-WPRE-HGH, and AAV5-Syn-FLEX-rc[ChrimsonR-tdTomato] (red). Data represented as mean ± S.E.M. **** $p < 0.0001$.

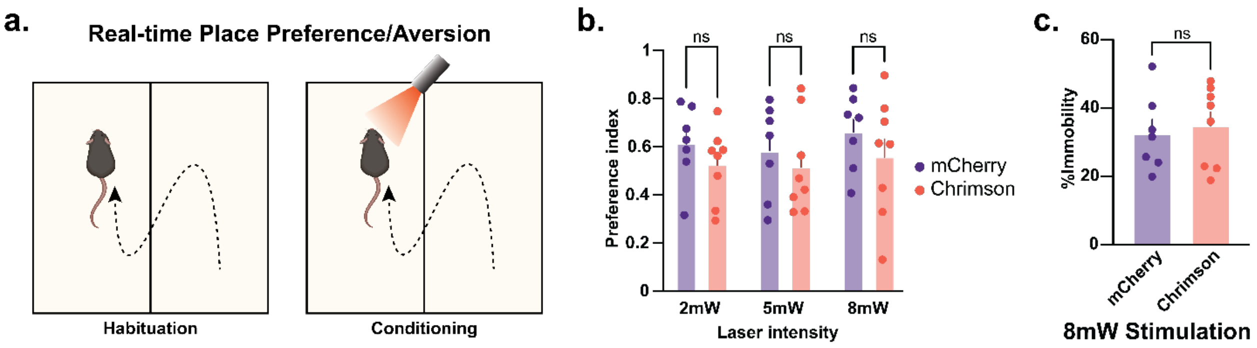


**Extended Data Figure 6. Optogenetic stimulation of DLS cholinergic interneurons does not produce place preference or place aversion.**

**(a)** Chrimson and mCherry control mice underwent CPP/CPA with optogenetic stimulation of DLS ChAT neurons occurring in either left or right chamber, counterbalanced.

**(b)** No significant difference in preference was seen between Chrimson and mCherry control mice at 2, 5, and 8mW (2-way ANOVA; $F(1, 13) = 1.254$, $p=0.2831$; Sidak post-hocs $p>0.05$; n=7-8 mice per group).

**(c)** No significant difference was seen in time spent immobile during optogenetic stimulation in CPP/CPA (Unpaired t-test; $t(13)=0.3793$, $p=0.7106$; Sidak post-hocs $p>0.05$; n=7-8 mice per group). Data represented as mean ± S.E.M. ns= not significant.

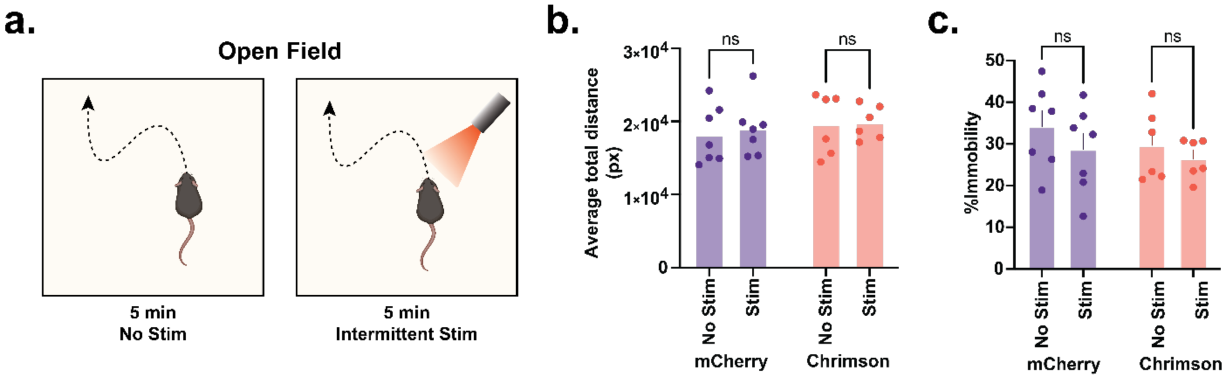


**Extended Data Figure 7. Optogenetic stimulation of DLS cholinergic interneurons does not alter locomotion or immobility in the open field.**

**(a)** Chrimson and mCherry control mice underwent an open-field task, where animals were allowed to explore freely for two 5 minutes trials (first 5 minutes without stimulation and second 5 minutes DLS ChAT neurons were optogenetically stimulated intermittently).

**(b)** No significant difference in distance travelled was seen between Chrimson and mCherry control mice with and without optogenetic stimulation (2-way ANOVA; Main effect of group: $F(1, 11) = 0.4593$, $p$=0.5120; Sidak post-hocs $p$>0.05; n=6-7 mice per group).

**(c)** No significant difference in time spent immobile was seen between Chrimson and mCherry control mice with and without optogenetic stimulation (2-way ANOVA; Main effect of group: $F(1, 11) = 0.5835$, $p$=0.41610; Sidak post-hocs $p$>0.05; n=6-7 mice per group). Data represented as mean ± S.E.M. ns= not significant.

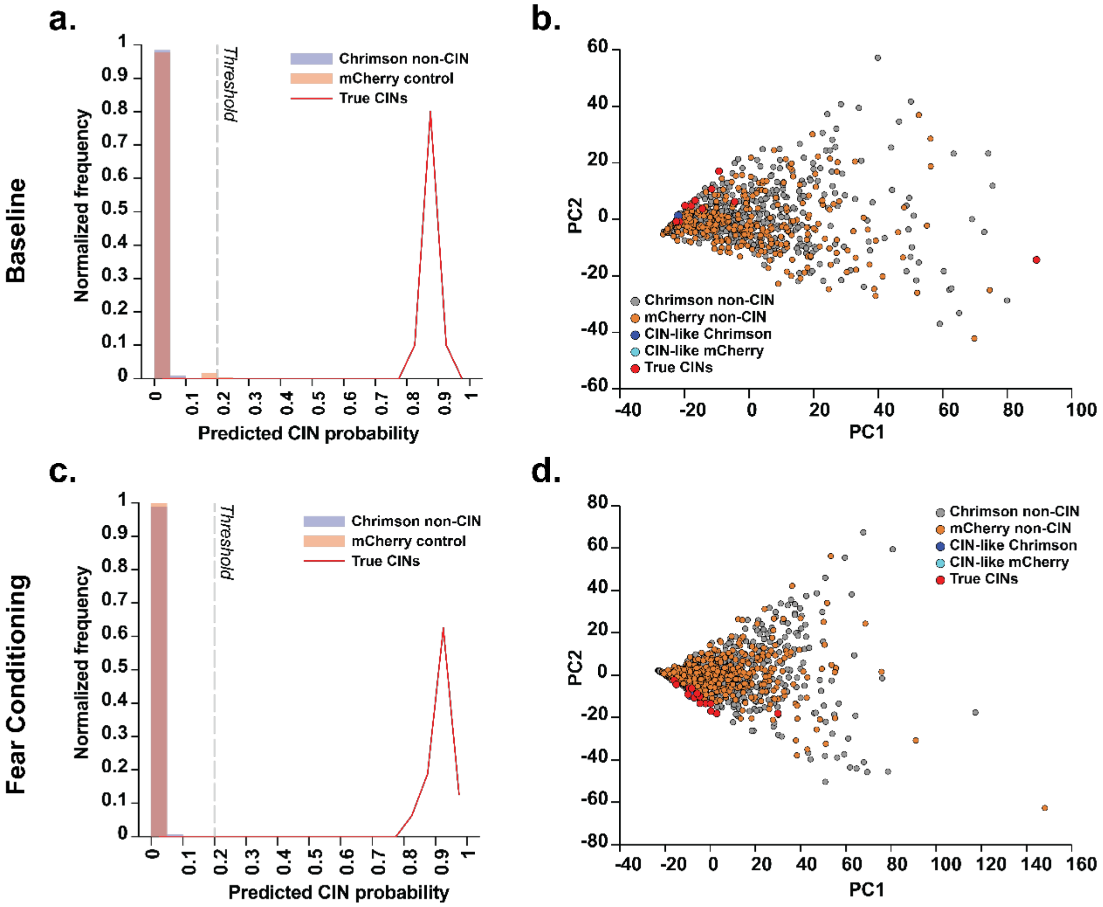


**Extended Data Figure 8. Random-forest classifier–derived CIN probability distributions and PCA clustering for baseline and fear-conditioning datasets.**

**(a–b)** CIN-probability distributions and PCA feature-space projections generated using a random-forest classifier for neurons recorded during baseline sessions with randomly delivered LED pulses. The classifier identifies true CINs (red) as a distinct high-probability peak, and PCA shows these neurons forming a tight, compact cluster in feature space. Nearly all other neurons in both Chrimson and mCherry control mice exhibit near-zero CIN probability and occupy non-CIN regions of the PCA space. Only a very small subset of neurons displays elevated CIN-like probabilities (blue for Chrimson, cyan for mCherry).

**(c–d)** Application of the same random-forest classifier to neurons recorded during fear-conditioning sessions yields a similar structure: true CINs again cluster separately with high probability, while the majority of neurons display low CIN probability and are distributed broadly across PCA space. CIN-like neurons remain rare in both groups.

Together, these analyses confirm that the random-forest classifier reliably distinguishes true CINs from the overwhelmingly non-cholinergic population across both baseline and fear-conditioning datasets, with CIN-like neurons representing only a minimal, well-defined minority.

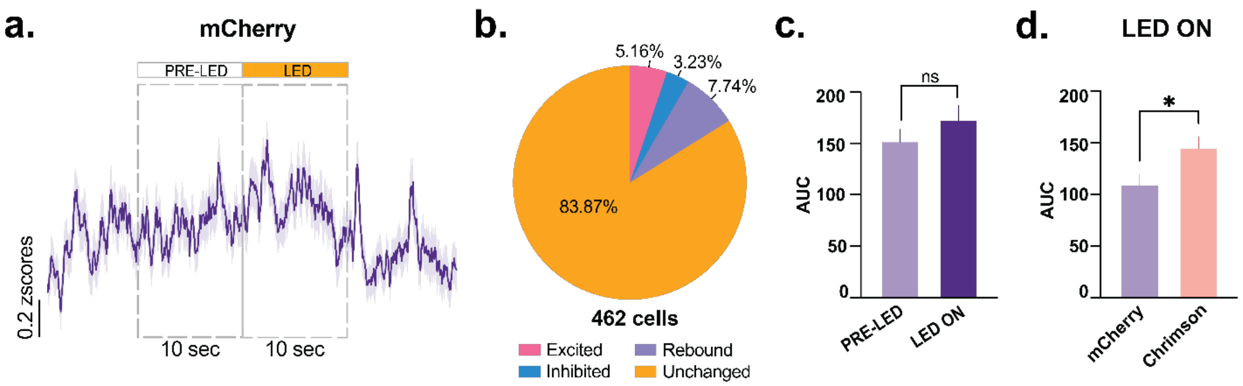


**Extended Data Figure 9. Comparison of baseline and LED-evoked DLS calcium activity in mCherry control versus Chrimson mice.**

**(a)** Baseline DLS calcium activity in mCherry control mice before and during LED stimulation.

**(b)** Proportions of DLS neurons showing excitation, inhibition, rebound responses, or no change during LED stimulation in mCherry control mice.

**(c)** No significant difference was seen in DLS calcium activity before and during LED stimulation (Paired t-test; $t(1243)=1.540$, $p=0.1237$; n=311 cells x 4 LED stims = 1244 LED stimulations).

**(d)** DLS calcium activity is significantly increased from baseline at the time of LED stimulation in Chrimson mice compared to mCherry controls (Welch's t-test; $t(3124)=2.307$, $p=0.0211$; n=1244-2052 LED stimulations). Data represented as mean ± S.E.M. * $p < 0.05$, ns= not significant.

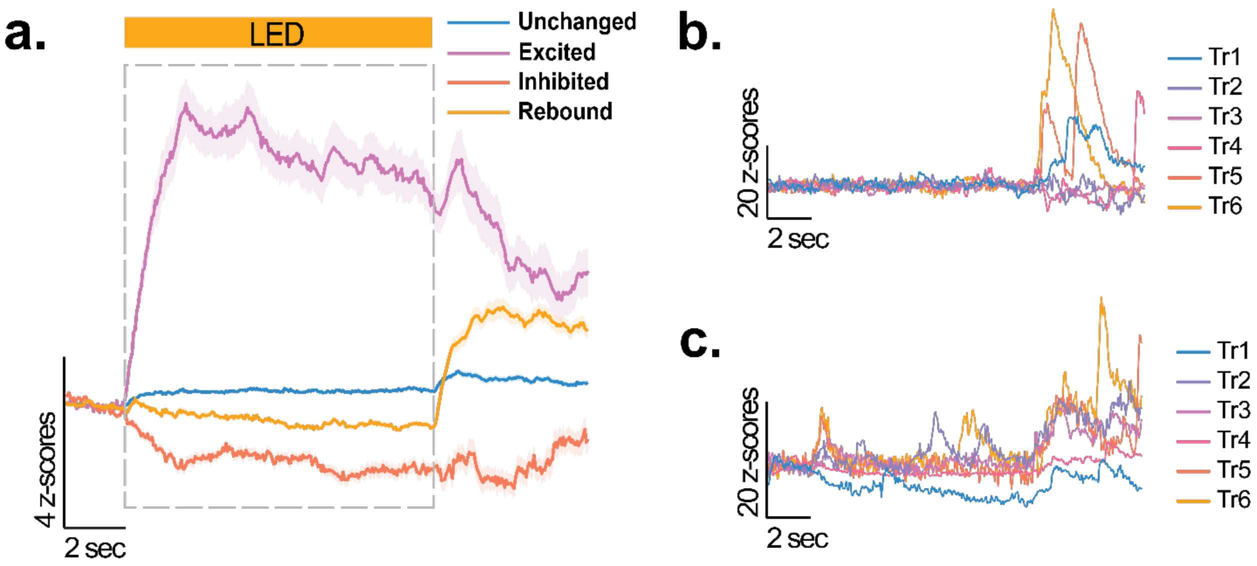


**Extended Data Figure 10. DLS calcium response profiles to LED stimulation and trial-by-trial activity during fear conditioning.**

**(a)** Mean DLS calcium activity in Chrimson mice at the time of LED stimulation, where neurons are either excited (purple), inhibited (orange), rebound (yellow), or are unchanged (blue).

**(b)** Representative calcium activity in fear conditioning during each trial for one Chrimson mouse.

**(c)** Representative calcium activity in fear conditioning during each trial for one mCherry mouse.

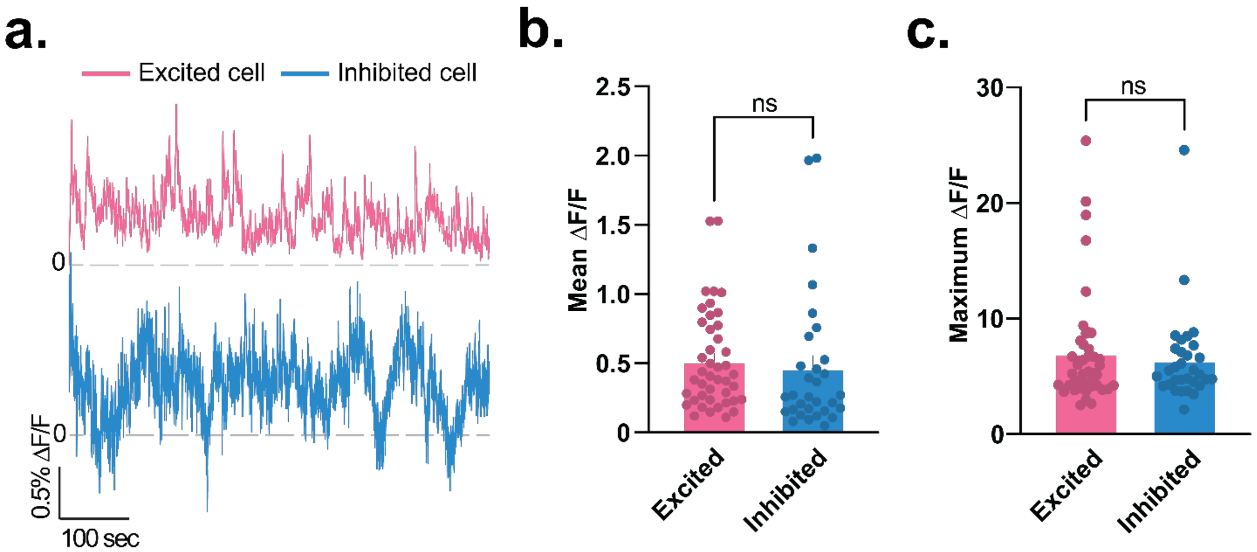


**Extended Data Figure 11. Baseline activity profiles of excited versus inhibited DLS neurons during fear conditioning.**

**(a)** Mean ΔF/F calcium traces (LED periods removed) from fear-conditioning sessions for neurons classified as *excited* (pink) or *inhibited* (blue) during optogenetic CIN stimulation.

**(b)** Excited and inhibited neurons do not differ in their mean ΔF/F amplitude across the session (Welch's t-test; $t(50.60)=0.5107$, $p=0.6118$; n= 31-43 cells per group).

**(c)** Maximum ΔF/F event amplitudes are also comparable between excited and inhibited neurons (Welch's t-test; $t(70.56)=0.5709$, $p=0.5698$; n= 31-43 cells per group). Together, these results suggest that the differences in inhibitory versus excitatory responses to CIN stimulation are not attributed to baseline calcium activity variations. Data represented as mean ± S.E.M. ns= not significant.

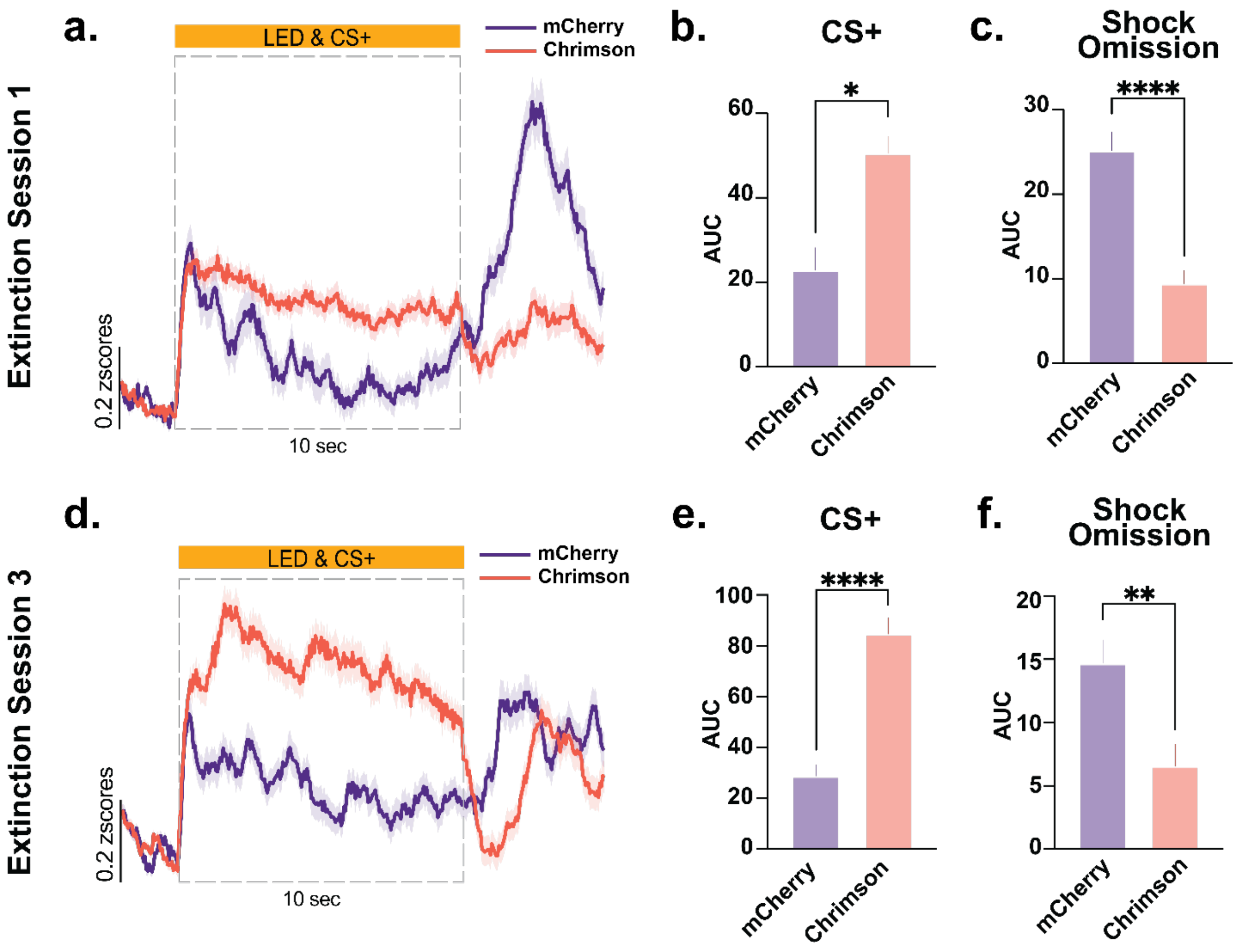


**Extended Data Figure 12. DLS population calcium responses to CS$^+$ and shock omission during extinction learning in mCherry control and Chrimson mice.**

**(a) Extinction Session 1:** Comparison of DLS ensemble activity between mCherry controls and Chrimson mice during the early phase of extinction. Mean ΔF/F traces during the CS+ and shock omission epochs reveal larger CS+-evoked responses in Chrimson mice relative to mCherry controls.

**(b)** CS+-evoked AUC is significantly higher in Chrimson mice compared to mCherry control mice during Extinction 1 (Welch's t-test; $t(1085)=2.352$, $p=0.0188$; n= 371-789 cells per group).

**(c)** Shock omission responses are significantly reduced in Chrimson mice relative to mCherry controls (Welch's t-test; $t(885.3)=4.587$, $p<0.0001$; n= 371-789 cells per group).

**(d) Extinction Session 3:** Comparison of DLS ensemble activity between groups during late extinction. Mean ΔF/F traces during the CS$^+$ and shock omission epochs show that Chrimson mice continue to exhibit elevated CS+-evoked responses compared to mCherry controls.

**(e)** CS+-evoked DLS neural ensemble calcium response remains significantly higher in Chrimson mice during Extinction 3, indicating persistence of CS+-related neural activity with CIN stimulation (Welch's t-test; $t(630.1)=3.345$, $p=0.0009$; n= 394-518 cells per group).

**(f)** Shock omission responses remain significantly lower in Chrimson mice compared to mCherry control mice during Extinction 3 (Welch's t-test; $t(901.4)=2.230$, $p=0.0260$; n= 394-518 cells per group). Together, these results show that DLS ACh release enhances and sustains

CS+-evoked ensemble activity across extinction; at the same time, shock omission responses remain suppressed, indicating a selective persistence of cue-related threat expectancy with CIN stimulation. Data represented as mean ± S.E.M. * $p < 0.05$, ** $p < 0.01$, **** $p < 0.0001$, ns= not significant.
